# Within-species variation eclipses between-species differences in *Pan* consolation

**DOI:** 10.1101/2024.07.17.601006

**Authors:** Jake S. Brooker, Christine E. Webb, Stephanie Kordon, Frans B. M. de Waal, Zanna Clay

**Affiliations:** Department of Psychology, Durham University, South Road, Durham, DH1 3LE, UK; Department of Human Evolutionary Biology, Harvard University, 11 Divinity Avenue, Cambridge, MA 02138, USA; Department of Psychology, Emory University, 36 Eagle Row, Atlanta, GA 30322, USA

**Keywords:** bonobo, chimpanzee, consolation, empathy, *Pan*

## Abstract

Empathy and its subcomponents are well documented throughout the animal kingdom, indicating the deep evolutionary origins of this socioemotional capacity. A key behavioural marker of empathy is consolation, or unsolicited bystander affiliation directed towards distressed others. Consolation has been observed in our closest living relatives, bonobos (*Pan paniscus*) and chimpanzees (*P. troglodytes*). However, systematic comparisons are absent, despite potential for interspecific differences. Bonobos are often considered less aggressive, more emotionally sensitive, and more socially tolerant than chimpanzees—key characteristics purported to drive consolation. Furthermore, social and individual factors also appear to drive intraspecific variation in empathy. To address within- and between-species variability in *Pan* consolation, we systematically tested the consolatory tendencies of large sanctuary-living groups of both species. Bonobos and chimpanzees exhibited similar consolation tendencies; however, within-species analyses revealed further similarities and variation. Bonobo consolation was most often directed towards and received by younger individuals, while chimpanzee consolation was most often directed towards close social partners. In addition, males and females of both species showed decreased consolation with age, with some evidence for chimpanzee males consoling more than young females. Our findings support the notion that within-species variation in *Pan* socio-emotional abilities is greater than between-species differences, highlighting the presence of striking behavioural diversity across our two closest cousins.

## Introduction

Empathy, broadly defined as sharing and/or understanding others’ emotional states, is a cornerstone of the human experience [1,2]. From politics and marketing to day-to-day social interactions, empathy allows us to effectively cooperate and communicate as well as strengthen our social relationships [3]. Cross-cultural and personality studies have revealed remarkable individual and cultural variation in empathic concern and perspective-taking [4–7]. Some longitudinal studies indicate that individual differences in human empathic responding are relatively stable [5–7]. As variation has been reported on individual and cultural levels, it has been suggested that both personality and cultural norms may influence empathy. Likewise, it has often been suggested that humans exhibit a uniquely advanced capacity for empathy, related to our advanced socio-cognitive capacities including perspective taking, self-other differentiation, emotion regulation and cognitive appraisal [8]. However, comparative research has revealed that empathy likely manifests through bottom-up processes from foundational components with a potential ancient ancestry in our evolutionary lineage [9]. These processes are thus proposed to be regulated by sophisticated top-down information processing mechanisms.

In the animal kingdom, many behaviours possibly related to empathy have been reported, mainly in mammals [10,11] though there is also some evidence in birds (e.g., consolation in *Corvus corax*; [12]). Empathy-driven behaviours emerge in the form of sub-components that represent differing expressions of emotional responsiveness and understanding [3,13]. Some of these sub-components, such as emotional contagion, have been observed across many social mammals, indicating a deep evolutionary history [10,13]. Furthermore, some behavioural prosocial manifestations of empathy, such as consoling others in distress, have been observed in rodents [14], elephants [15], and some primate and great ape species (for a review, see [13]).

Consolation, defined as unsolicited affiliation offered by an uninvolved bystander towards a distressed conspecific [16], is often considered a more complex empathic behaviour. Consolation, sometimes referred to as ‘sympathetic concern’ [17], is thought to require both a cognitive appreciation (and even understanding) of another’s state combined with a prosocial orientation to improve it, such as by providing reassuring contact [18]. Among non-human animals, our two closest relatives, the bonobos and chimpanzees have demonstrated the capacity for consolation in a range of settings, including captivity, sanctuaries, and even the wild [19–23]. Among wild chimpanzees, both the western and eastern subspecies have been observed to console others, including at Taï National Park, Ivory Coast [22] and the Mahale Mountains National Park, Tanzania [24] respectively. Another study of the eastern subspecies in Budongo Forest, Uganda, showed that consolation only occurred after 3.3% of observed conflicts, which was not significantly more than matched controls [25]. In contrast, Kutsukake and Castles [24] observed consolation after around 22% of conflicts in the Mahale M-group of eastern chimpanzees. This difference between the Mahale and Budongo groups indicate possible within-species behavioural flexibility and cross-community variation in consolatory tendencies. Whilst there are no studies of consolation in wild bonobos, case studies of targeted prosocial behaviour, such as conspecifics attempting to remove snares from group members and searching for lost group members [26] suggest consolation is likely. Moreover, consolation has been observed in sanctuary-settings [19], including by wild-born immature individuals, suggesting that consolation is likely part of the wild bonobo repertoire.

Evidence that immature apes engage in consolation challenges assumptions that consolation is dependent on sophisticated cognitive mechanisms. In both chimpanzees and bonobos, consolation is offered most by younger individuals [19,27], suggesting that empathic orientation towards other’s states may be intrinsic and adaptive. In humans, concern for others including consolation has been reported even within the first year of life, as early as nine months [28]. Empathic behaviours, like consolation, may strengthen interpersonal bonds, especially for dyads that demonstrate enduring cooperative relationships. For this reason, social closeness, familiarity, or similarity has been shown to be a key predictor of empathic tendencies in humans [29,30] and among nonhuman apes (bonobos: [19,31]; chimpanzees: [21,27,32]).

Bonobos and chimpanzees develop long term enduring bonds with other group members and form strong support networks [33]; yet there are reported differences in their social tendencies and socio-emotional orientations. For example, bonobos appear to have enhanced attentiveness to conspecific social and emotional expressions than chimpanzees [34,35]. Apparent differences in sociability, socio-sexuality, and aggressive tendencies between the two species may have biological and neurological foundations. Firstly, chimpanzees, but not bonobos, have been shown to have deletion of the DupB region in the AVPR1A gene, which includes a microsatellite called RS3 [36]. Variation in RS3 is linked with variation in social bonding, and increased levels in bonobos may support their reported xenophilia. Neurological research shows that bonobos have twice the density of serotonergic axons in the amygdala than chimpanzees [37]. This region is associated with social cognition and emotional regulation amongst others, and this variation between species may mediate the differences in their social structures and behaviour. The ability to regulate one’s own emotions—i.e., suppressing personal arousal upon exposure to an arousing situation—can vary between individuals [38]. However, emotional regulation appears to be crucial for empathy to emerge [39]. In addition, bonobos have more grey matter in the right dorsal amygdala and right anterior insula than chimpanzees, as well as a larger pathway linking the amygdala with the ventral anterior cingulate cortex [40]. These regions are associated with perceiving distress in oneself and others, as well as control of reactive and proactive aggressive impulses. These differences, as well as lower purported levels of aggression in bonobos, and therefore decreased risk of injury [41,42], may facilitate enhanced production of prosocial empathic behaviours, such as consolation and targeted helping, compared to chimpanzees.

Social tolerance is also thought to influence empathic tendencies, according to the *Social Constraints Hypothesis* [43]. For example, evidence of consolation in the Tonkean macaques (*Macaca tonkeana*; [44]) and not in other more despotic macaque species (for a review, see [13]), indicates that social tolerance may facilitate empathic responses that might otherwise be inhibited due to intimidation, fear, or aggression. Chimpanzees are often considered to be more intolerant and despotic than bonobos [33], following evidence that chimpanzees have formal signals of subordinacy, through submissive gestures [45], whereas bonobos do not [46]. These expressions of submission are often used to assess chimpanzee hierarchies, which commonly follow a linear pattern [47–49]. Bonobos appear to have a more flexible, egalitarian dominance style [50–52], which may be facilitated by their reduced aggressivity and increased sociability. Furthermore, chimpanzees tend to be xenophobic to other communities [53]. In contrast, whilst wild bonobo communities can be vocally hostile to other groups, numerous populations regularly have peaceful intergroup encounters, even cases of food sharing, and exchanges of sex, grooming, and play [54,55]. Some experimental paradigms have indicated that bonobos outperform chimpanzees on cooperation [56] and theory-of-mind-related tasks [57]. Where empathy expressions are related to understanding others, emotions, and social tolerance, one might expect bonobos to be more consolatory than chimpanzees.

However, there is evidence of within-species variation. Several semi-wild chimpanzee communities have been shown to vary in interindividual tolerance measured by cofeeding proximity [58]. Similarly, a recent comparison of multiple captive and semi-wild populations revealed that levels of social tolerance overlapped between bonobos and chimpanzees [59]. Furthermore, as previously mentioned, consolation appeared relatively absent in one population of eastern chimpanzees at Budongo [25] and yet present in their Mahale counterparts [24]. These communities have been demonstrated to vary considerably in social tolerance and hierarchical steepness, whereby Budongo chimpanzees have steeper, and more despotic hierarchies compared to Mahale [60]. Whilst these wild groups have not been systematically compared, these findings indicate that consolation may show inter-group variation and emerge flexibly if conditions permit. As population-level variation in group social tolerance can differ within species, consolatory tendencies may vary across groups living in similar conditions.

Whilst some socio-cognitive abilities of bonobos and chimpanzees have been compared, direct systematic comparisons of their consolation tendencies have yet to occur. As coalitionary support and intermediate despotism are key factors of bonobo and chimpanzee societies, comparing them may reveal whether reduced physical aggression and increased emotional orientation in bonobos predict an increased likelihood to console. Consolation does appear to be a feature of natural *Pan* social living; however, wild studies are scarce due to methodological constraints. Therefore, comparing great apes in semi-wild sanctuary settings can provide a balance of natural ecological surroundings with improved observational conditions. Based on evidence that bonobos show enhanced social awareness and emotional sensitivity, we tested the hypothesis that bonobos are more empathic than chimpanzees, predicting that consolation would be more prevalent in bonobos than chimpanzees. Furthermore, following the hypothesis that empathy is socially-biased [2], we predicted higher rates of consolation between dyads that are socially bonded, through kinship and close affiliative tendencies.

Consolatory tendencies tend to decrease in each species with age [19,27], whilst previous studies indicate a lack of general sex differences in consolation in bonobos and chimpanzees. However, as males remain in their natal groups in each species [46,47], they might be expected to invest in building long-term social bonds already at a young age. In bonobos, males tend not to have strong social bonds beyond their mothers. They have been shown to form bonds with other females that extend their alliance relationships and improve reproductive success, however these are typically females with elevated rank positions, therefore reducing the likelihood that they would be victims of conflict [61,62]. In chimpanzees, adult males are less likely to be victims of conflicts than females and immature males, implying that their closest social partners, other adult males, will have fewer opportunities to offer consolation. As such, and in line with previous findings [19,27], we predicted that, regardless of sex, younger individuals of both species will show the highest tendencies to console. In addition, we predicted an interaction between bystander age and sex in both species, whereby younger individuals of both sexes would show the highest consolatory tendencies whereas in adulthood, older males of each species would show lower rates than females. By comparing between- and within-species influences on an empathic behaviour like consolation in our closest living relatives, we aim to improve our understanding of the origins of human empathy and emotional responsiveness.

## Methods

### Subjects and housing

We conducted observations of bonobos at Lola ya Bonobo Sanctuary (hereafter, “Lola ya Bonobo”) in the Democratic Republic of the Congo during July-September 2019. We conducted observations of chimpanzees at Chimfunshi Wildlife Orphanage Trust (hereafter, “Chimfunshi”) in the Copperbelt Province of Zambia during March-August 2019. Lola ya Bonobo houses three groups of bonobos in enclosures ranging from 15–20 hectares with rainforest, lake, swamp, stream, and open grass area, whilst Chimfunshi is home to four groups of chimpanzees that are accessible for observational research, living in outdoor miombo woodland enclosures ranging from 20–80 hectares. At Lola ya Bonobo, bonobos sleep together in dormitories. At Chimfunshi, chimpanzees nest independently unless kept inside for monitoring or medical intervention.

Both sanctuaries house wild-born individuals, orphaned and rescued from the pet and bushmeat trades, as well as sanctuary-born individuals, all of whom are supported by caregiving and veterinary staff. In both sanctuary environments, the apes could roam, forage, and nest independently, whilst supported by an onsite caregiving team. The groups at Lola ya Bonobo sleep at night in a managed indoor dormitory and otherwise roam in their forested enclosures during the daytime. At both sites, groups are provisioned at least twice per day with a variety of fruits and vegetables.

At Lola ya Bonobo, we observed all inhabitants of Groups 1 and 2 (hereafter: B1 and B2), which housed *N* = 22 and *N* = 18 bonobos, respectively. At Chimfunshi we observed all inhabitants of Group 2 (hereafter: C2), the largest group which comprised *N* = 50 chimpanzees at the start of data collection. The age and sex composition of our sample is provided in Table S1 (ESM). We logged 800 hours of observations at Chimfunshi and 600 hours of observations at Lola ya Bonobo.

### Data collection

#### Victim focal follows

Consolation has typically been recorded using the post-conflict/matched-control method [63]. This involves conducting focal follows of a victim for a standardised period—usually 5–10-minutes—after a conflict or spontaneous distress period, while recording the initiator and behaviour of all affiliative interactions that occur involving the focal victim. Post-conflict (PC) and post-distress (PD) periods are then compared with matched control (MC) recordings observed at a similar time and circumstance a day later. The PC/MC method has already reliably demonstrated consolation in multiple bonobo and chimpanzee communities compared with matched controls [19–21,43], including in some of these sample populations [19]. Thus, we decided to only collect post-conflict and post-distress events to ensure a large enough sample to compare individual and social influences on consolation tendencies.

We used focal all-occurrence sampling [64] to collect *N* = 150 PCs and *N* = 10 PDs in C2. In B1, we observed *N* = 28 PCs and *N* = 36 PDs and in B2, we observed *N* = 36 PCs and *N* = 16 PDs. Our total sample of events was therefore *N* = 276 across both species and all three groups. Previous studies of consolation show that most consolatory interactions occur during the first minute after distress [19]. We therefore conducted victim focals for the first 5-minutes following distress to optimise our limited observation time. We defined agonistic encounters by the presence of at least one of the following behaviours: high-contact aggression [hit, slap, kick, trample, bite], low-contact aggression [poke, push, push away, brush aside], chase, or threat [threat bark, swagger, display, flail arm, stamp] [45]. PCs were only analysed if the victim elicited a clear victim response, identified as the occurrence of bared-teeth scream, whimper, tantrum, or flee from aggression [45] following an agonistic encounter. If these victim response behaviours occurred spontaneously, they were coded as PD events. During the 5-min focal-follows, we noted the presence of all bystanders within 1-metre and 5-metres of the focal subject at the onset of the event. We recorded all PC/PD victim focals using Panasonic HC-V770 Camcorders with detachable Sennheiser MKE 400 directional shotgun microphones.

#### Dyadic social affinity and gregariousness

We also collected focal scan follows [64] of social tendencies to measure dyadic affiliation levels and individual gregariousness. We collected social focals outside of feeding times for both species. As we measured affiliation on a dyadic level, we accumulated information on individuals who were not currently the focal subject if they associated with the focal. The data set consisted of *N* = 706 focal follows for B1 (*range* = 28–41; *M* = 32.1; *SD* = 11.0), *N* = 587 focal follows for B2 (*range* = 16–44; *M* = 34.2; *SD* = 11.3), and *N* = 684 focal follows for the C2 (*range* = 9–18; *M* = 14.3; *SD* = 2.6). Each focal follow lasted 10-minutes and behaviours were recorded at 1-minute scan points, upon which we recorded all social interactions the focal was involved in including grooming, play, and sex, as well as all individuals within 1-metre. As the social behaviours required proximity of 1-metre to the focal, we only coded either behaviour or proximity once per dyad per scan during observations. However, when we compared dyadic scores for behaviour and proximity when independent of one another, they were strongly correlated (r(375) = .43, *P* < .001). We thus incorporated all social behaviour observations into one score for proximity and used that to compute unique dyadic affiliation scores for every possible dyad. We divided the number of scan points each dyad interacted for by the total number of scan points both individuals of the dyad was observed for. We also computed individual gregariousness scores by dividing the total number of scans each focal subject was observed within proximity to at least one other individual by the total numbers of scans they were observed for. At Chimfunshi Wildlife Orphanage, we recorded social focal follows using the tablet app Zoo Monitor [65], which contributed towards long-term data collection of social relationships at this site. At Lola ya Bonobo, we recorded social focals by hand due to software incompatibility. In addition, we were interested in the effect of kinship, and defined each dyad as ‘kin’ or ‘non-kin’ depending on whether they shared a maternal genetic relationship or not, respectively. This therefore included all mother-infant, sibling-sibling, and grandparent-grandchild pairs as ‘kin’. Some dyads at Chimfunshi represented a genetic uncle-nephew dynamic and were also included as ‘kin’. Inter-observer reliability revealed almost perfect agreement between observers (partner identity: bonobo κ > 0.80; chimpanzee κ > 0.96; social behaviour: bonobo κ > 0.95; chimpanzee κ > 0.93).

### Hierarchies

We assigned categorical rank values (high, medium, low) for each ape. For both species, we used the R package ‘EloRating’ [66] to create dominance scores based on dyadic agonistic interactions involving high-contact aggression, low-contact aggression, and chases [66]. Each agonism type was assigned a different optimised weighting, known as a K-value, based on intensity and likelihood of winning probabilities. We only included individuals four-years and older with at least six observed dyadic agonistic interactions in this ELO analysis. For the chimpanzees, the ordered rank output was consistent with categorical rankings acquired from the keepers and care staff. Hence, we evenly divided this output into ‘high’, ‘medium’, and ‘low’ rank, and used the keeper allocations to assign categorical ranks to individuals for whom we lacked sufficient ELO data for. For the bonobos, the ELO analysis revealed ordered results that were broadly but not wholly consistent with keeper and researcher observations. Whilst human judgements of social rank may be influenced by observer bias, these slight inconsistencies were likely contributed to by a relatively lower quantity of agonistic observations over our relatively short study period, therefore affording our ELO analysis lower power. However, whilst the rank order was inconsistent, categorisation into ‘high’, ‘medium’, and ‘low’ was fairly consistent with keeper and researcher observations. As we did not have substantial or prior quantitative data to support the ELO analysis, we decided to assign rank categories based on agreed deliberation between two long-term experienced observers of the bonobos at Lola ya Bonobo, supported by views of the sanctuary staff. For both species, we assigned mother ranks to infants (apes between 0 and 2 years of age). In both cases, observer rankings were based on observations of dynamics surrounding food and resource competition.

### Ethics

This study was approved by the Senior Veterinary Advisory Team of Lola ya Bonobo Sanctuary, the Chimfunshi Research Advisory Board, and the Animal Welfare Ethics and Research Board of Durham University. Data collection comprised purely naturalistic observations and adhered to the legal requirements of the Democratic Republic of the Congo and Zambia, as well as the International Primatological Society’s Principles for the Ethical Treatment of Nonhuman Primates.

## Analysis

We conducted all-occurrence coding of affiliative and agonistic encounters during PC/PD periods in ELAN [67,68]. Consolation was defined as the onset of an affiliative interaction involving contact during the 5-minute follow that was spontaneously initiated by a bystander. Initiation was defined as the solicitation of an affiliative interaction either by gesturing or initiating physical contact with a partner. Thus, interactions initiated by the victim—e.g., the victim approached the bystander—were excluded as they do not represent spontaneous approaches by the bystander, thereby these interactions are not considered consolation [69]. Affiliative contact behaviours included body kiss, contact sit, embrace, finger/hand in mouth, embrace, genital inspection, genito-genital contact, grasp hand, groom, hunch-over, mount, mount walk, mouth kiss, pat, play, rump-rump touch, and touch [45,70]; see Table S2). Inter-coder reliability of consolation occurrence and consoler identity indicated almost perfect agreement (consolation occurrence: bonobo κ = 0.86, chimpanzee κ = 0.85; consoler identity: bonobo κ = 1.00, chimpanzee κ = 0.97).

### Summary of statistical approaches

We used a Bayesian mixed models approach to assess whether consolation was more likely to occur in the bonobo or chimpanzee groups sampled, along with within-species variation in consolation tendencies according to bystander and victim characteristics: sex, age, rank, gregariousness, and group, as well as the bystander-victim social relationship. We collapsed our observations into two formats: event-level, where each observation row represents one PC or PD event; and dyad-level, where each observation row represents a unique bystander-victim combination within an event. This allowed us to comparably investigate general species tendencies at the event-level and then to assess the influence of individual (e.g., bystander age, bystander sex) and social factors (e.g., kinship, affiliation score) on consolation at the dyad-level. We z-transformed all covariate predictors and dummy coded and centred all factor predictors prior to their inclusion as random slopes.

We fitted all models in RStudio (version 1.3.1093; [71]) using the function brm of the package brms (version 2.21.0; [72]). Each model included four Markov chain Monte Carlo (MCMC) chains, with 10,000 iterations per chain, of which we specified 2,000 iterations as warm-up to ensure sampling calibration. This resulted in 32,000 posterior samples in total for each model. In all models, we used default priors (weakly informative priors with a student’s t-distribution of 3 degrees of freedom and a scale parameter of 2.5) to ensure that our inferences were driven primarily by the data. Flat priors cover a wide range of parameter values without favouring any specific region. This allowed our already complex models incorporating many fixed and random effects to explore a broad parameter space, ensuring that no potential values are excluded a priori. Furthermore, flat priors emphasize the likelihood function, allowing the posterior distribution to closely follow the shape of the likelihood. Lastly, this approach also facilitates comparison with frequentist methods, enhancing the interpretability and generalisability of our findings.

For all models, we used multiple measures to summarise the posterior distributions for each variable and report evidence of an effect: 1) we characterised uncertainty by two-sided credible intervals (89% CrI) [73], denoting the range of probable values in which the true value could fall [74]; and 2) we computed the probability of direction (hereafter: *pd*) [75]. The 89% CrI reveals the range within which an effect falls with 89% probability [73], while the *pd* indicates the probability that a parameter is strictly positive or negative [75]. For all models, diagnostics revealed an accurate reflection of the original response values by the posterior distributions, as R-hat statistics were < 1.01, the numbers of effective samples >1000, and MCMC chains had no divergent transitions (see electronic supplementary material, hereafter: ESM, Figure S1) [72]. Furthermore, diagnostics also revealed an accurate reflection of the original response values by the posterior distributions (see ESM, Figure S2).

### Event-level analyses: Testing species differences

We fitted two Bayesian Generalised Linear Mixed Models (GLMMs) to investigate whether consolation was more likely to occur in bonobos or chimpanzees. The sample for this model consisted of *N* = 276 events (PC *N* = 214; PD *N* = 62) of *N* = 90 apes (bonobo *N* = 40; chimpanzee *N* = 50). The first model (hereafter: *Model 1.1*) fitted a Bernoulli distribution (consolation occurrence: 1 = yes; 0 = no) and the second model (hereafter: *Model 1.2*) fitted a Poisson distribution (count of consolations per event). The fixed effects structure for these models included species and the number of bystanders. As population sizes varied across the three groups, and consolation may be more likely if there are more individuals present, we included the number of bystanders as a control effect to improve the accuracy of the estimated fixed effect of species. The random effects structure consisted of random intercepts for aggressor identity and victim identity, as well as random slopes for number of bystanders within both aggressor identity and victim identity.

### Dyad-level analyses: Testing individual and social factors

To test how individual and social factors influenced consolatory tendencies, we fitted two GLMMs to our dyad-level data for each species separately [76]. Each observation row represented a particular bystander-victim combination present during a given victim follow event. Two bonobos and three chimpanzees were excluded due to long periods of absence from their main groups (bonobos: 62–95% of observation time; chimpanzees: 54–65% of observation time). The sample for the bonobo model (hereafter: *Model 2.1*) consisted of *N* = 1656 observation rows from *N* = 38 individuals observed across *N* = 116 events within *N* = 2 groups. The sample for the chimpanzee model (hereafter: *Model 2.2*) consisted of *N* = 5668 observation rows from *N* = 47 individuals observed across *N* = 160 within *N* = 1 group.

Our response variable constituted a binary outcome fitted with a Bernoulli distribution according to whether the bystander solicited affiliative contact to the victim focal during the 5-minute event (consolation occurrence: 1 = yes; 0 = no). Full models included individual factors of the bystander (age, rank, sex), the victim (age, rank, and sex), and the social relationship of the dyad (kinship) as fixed effects. Unfortunately, due to inconsistencies in the social affiliation data collection protocol between Lola ya Bonobo and Chimfunshi, we were unable to include quantitative measures for gregariousness and dyadic affiliation score in the bonobo model. The chimpanzee model did however included fixed effects for bystander gregariousness and dyadic affiliation score. For both species, we also included the interaction of bystander age and bystander sex as a fixed effect and the bonobo model additionally included group as a fixed effect. The random effects structure for each model consisted of random intercepts for the individual identities of the aggressor, bystander, and victim as random effects as well as the event ID. We also included theoretically identifiable random slopes of victim age and sex within bystander identity and bystander age and sex within victim identity.

## Results

### Event-level results: Do bonobos console more than chimpanzees?

Across observations, consolation occurred at least once in 127 of the 276 events (46.0%). On average, *M* = 0.64 (*SD* = 0.81) consolatory approaches occurred per event in bonobos, and *M* = 0.75 (*SD* = 1.00) consolatory approaches per event in chimpanzees. *Model 1.1* revealed no effect of species on the probability of consolation occurring at least once in an event (*b* = 0.08, *SD* = 0.33, 89% credible interval (CrI) [−0.94, 1.11], *pd*: 0.55; see Figure 1). Additionally, *Model 1.2* showed no effect of species on the count of consolers that responded per event (*b* = 0.08, *SD* = 0.33, 89% CrI [−0.44, 0.60], *pd* = 0.60). Importantly, in each model, the inclusion of the number of bystanders also did not influence the outcome variables (*Model 1.1*: *b* = 0.13, *SD* = 0.15, 89% CrI [−0.36, 0.62], *pd* = 0.66; *Model 1.2*: *b* = 0.13, *SD* = 0.15, 89% CrI [−0.12, 0.37], *pd* = 0.80), indicating consolation likelihood does not increase if more possible consolers are present. Output from the event-level models can be seen in Table S3 (ESM).

**Figure 1.**
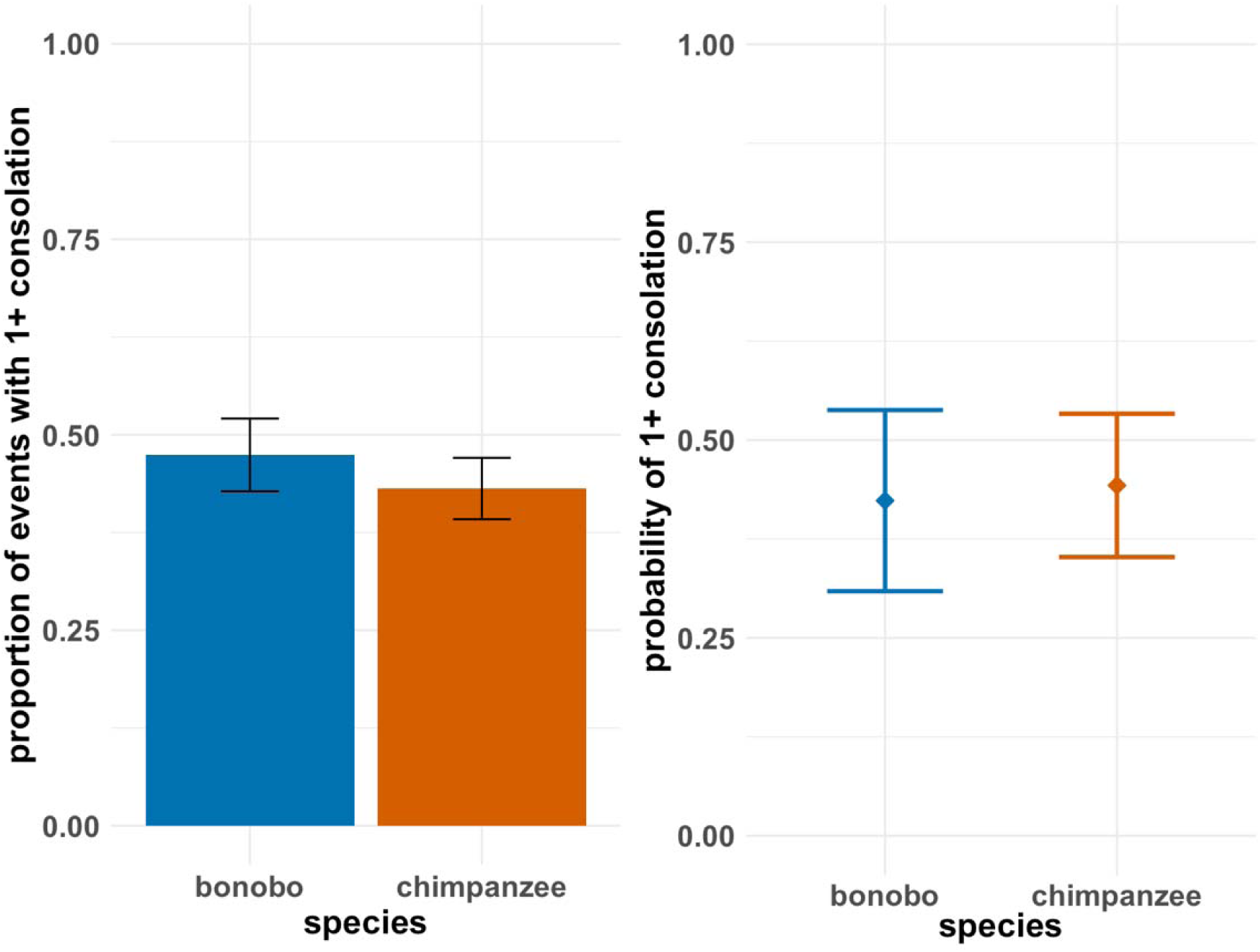
Barplot (left) and conditional probability plot (right) showing no credible difference between bonobos and chimpanzees in their relative tendency for a PC or PD event to feature 1+ consolation approach towards a distressed victim. Results obtained from *Model 1.1*. Whiskers on barplot show one standard error above and below mean. Probability plot shows conditional effect of species with 89% credible intervals.

### Dyad-level results: Which within-species factors drive consolation tendencies?

A full table of the output for bonobos (*Model 2.1*) and chimpanzees (*Model 2.2*) can be seen in Table S4 (ESM).

#### Social effects

We found a very strong effect of kin dyads consoling more than non-kin dyads in both species (bonobos: *b* = −2.35, *SD* = 0.62, 89% CI [−3.37, −1.37], *pd* = 1.00; chimpanzees: *b* = −0.86, *SD* = 0.41, 89% CrI [−1.50, −0.19], *pd* = 0.98; see Figure 2). We also found a very strong positive effect of affiliation level on consolation in chimpanzees (*b* = 0.20, *SD* = 0.09, 89% CrI [0.05, 0.35], *pd* = 0.98; see Figure 2). Furthermore, in chimpanzees, we found no clear effect of bystander gregariousness (chimpanzees: *b* = −0.16, *SD* = 0.21, 89% CrI [−0.49, 0.17], *pd* = 0.79). Finally, for bonobos we found no clear evidence for an effect of group (*b* = −0.49, *SD* = 0.52, 89% CrI [−1.34, 0.32], *pd* = 0.83).

**Figure 2.**
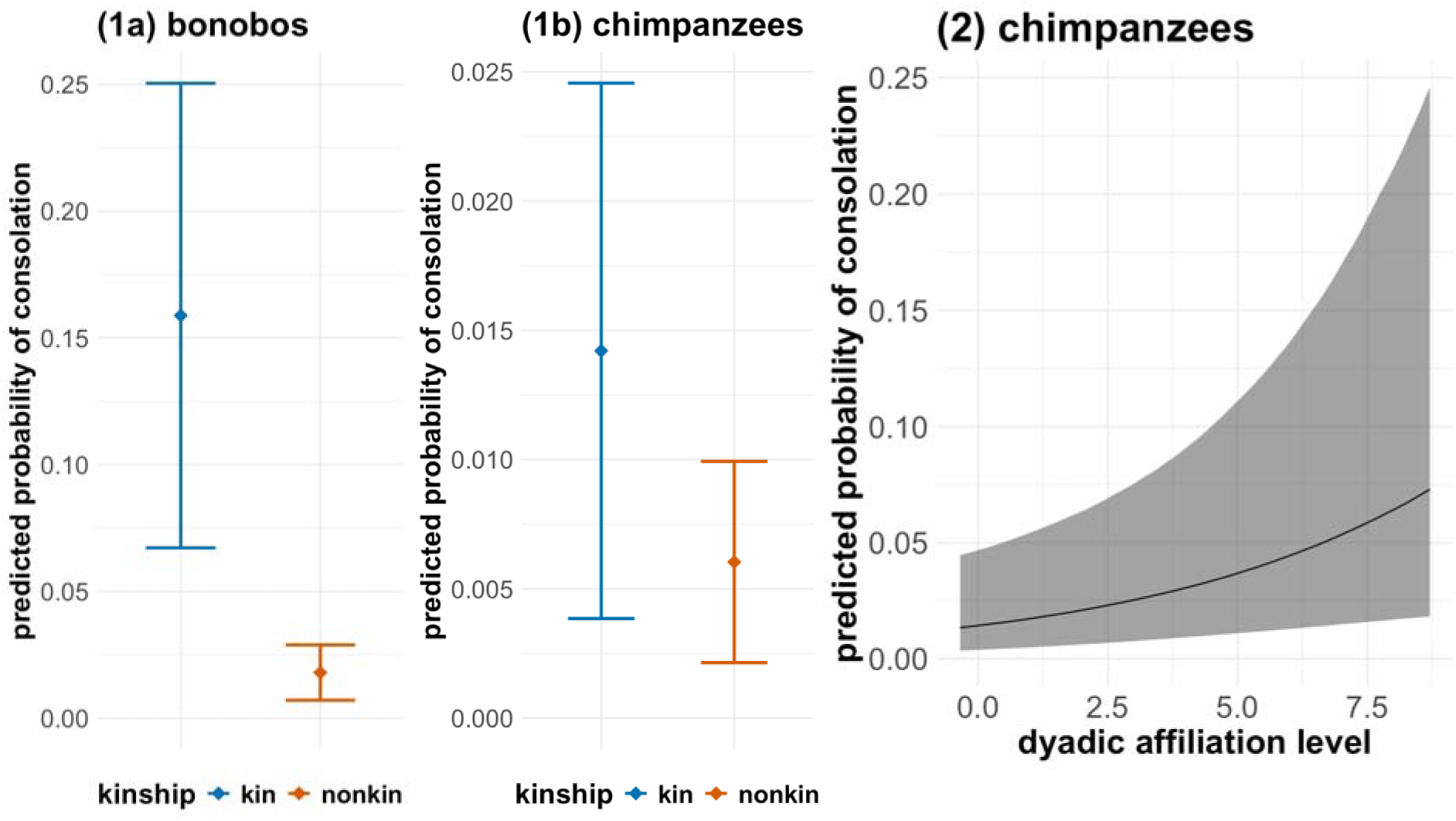
Predicted probability plots showing credible effect of kinship in bonobos (1a) and chimpanzees (1b) and a credible effect of dyadic affiliation level in chimpanzees (2). For all plots, Y-axis = predicted probability of consolation. (1a-1b) X-axis = kinship. Square points and error bars correspond to posterior means and the upper and lower 89% credible intervals, respectively. (2) X-axis = z-transformed dyadic affiliation level. Shaded area corresponds to 89% credibility intervals. Dyadic affiliation level not tested in bonobos.

#### Victim characteristics

In bonobos, we found a very strong negative effect of victim age where younger bonobos were consoled more than older bonobos (*b* = −1.59, *SD* = 0.74, 89% CrI [−2.88, −0.55], *pd* = 1.00; see Figure 2), but no credible effect in chimpanzees (*b* = 0.28, *SD* = 0.25, 89% CrI [−0.13, 0.68], *pd* = 0.87; see Figure 2). We found no credible effects for victim sex in either species (bonobos: *b* = 1.08, *SD* = 0.97, 89% CrI [−0.36, 2.68], *pd* = 0.88; chimpanzees: *b* = −0.56, *SD* = 0.65, 89% CrI [− 1.62, 0.41], *pd* = 0.81). Victim rank had no credible effect on consolation tendency in either bonobos or chimpanzees (see ESM, Table S4).

**Figure 3.**
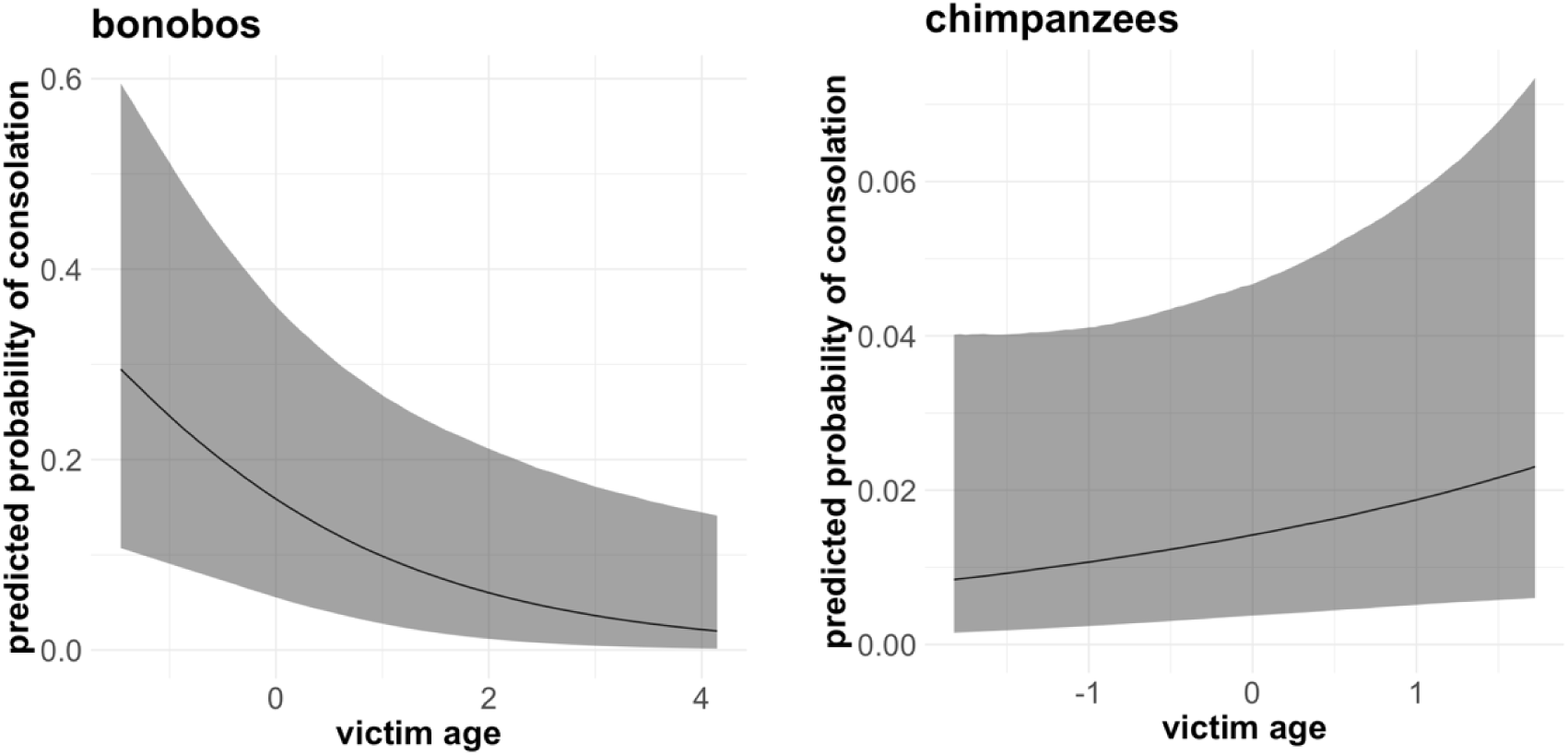
Predicted probability plot showing credible main effect of victim age in bonobos (left) and no credible effect in chimpanzees (right). X-axis = z-transformed age. Y-axis = predicted probability of consolation. Shaded area corresponds to 89% credibility intervals.

#### Bystander characteristics

We found no clear effect of bystander age and bystander sex interacting in bonobos (*b* = −0.81, *SD* = 0.89, 89% CrI [−2.32, 0.50], *pd* = 0.67), however, there was evidence of a strong positive effect of bystander age (*b* = −0.74, *SD* =, 89% CrI [−1.64, 0.12], *pd* = 1.00; see Figure 4) indicating that younger bystanders may console more than older bystanders. However, for chimpanzees we found moderate evidence for an interaction between bystander age and bystander sex (*b* = −0.76, *SD* = 0.61, 89% CrI [−1.77, 0.19], *pd* = 0.90; see Figure 4). This interaction indicates that indicated that younger individuals of both species consoled more than older individuals, with young males consoling more than females. However, males appear to decrease in their consolation tendency at a steeper rate than females. Conditional probability plots of these interactions are shown in Figure S3 (ESM). Bystander rank had no credible effect on consolation tendency in chimpanzees (see ESM, Table S4). However, high-ranking bonobos consoled victims more than low-ranking bonobos (low vs. high: *b* = −1.01, *SD* = 0.50, 89% CrI [−1.81, −0.24], *pd* = 0.98) but there was no credible difference between medium- and high-ranking bystanders (medium vs. high: *b* = −0.43, *SD* = 0.51, 89% CrI [−1.44, 0.57], *pd* = 0.81).

**Figure 4.**
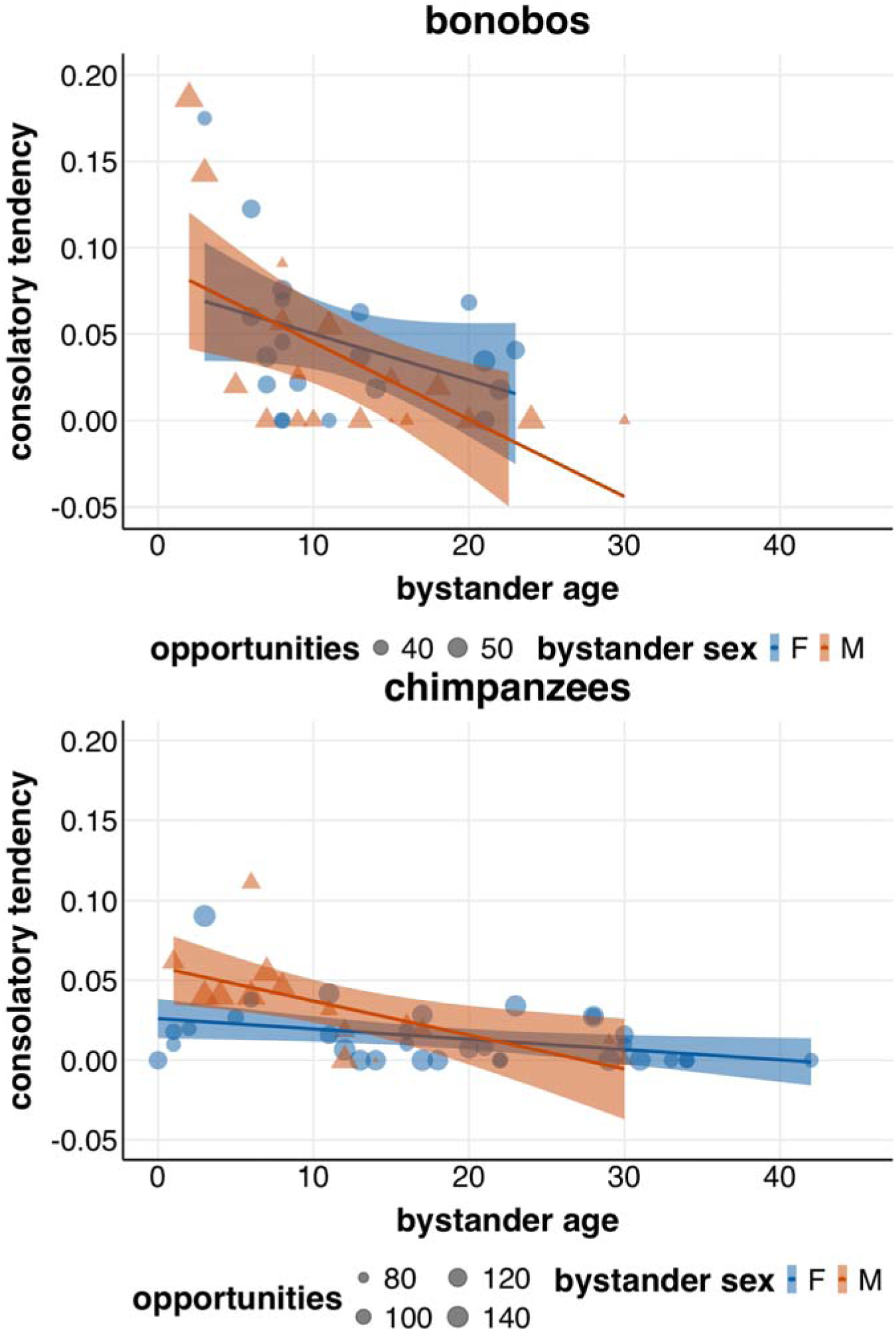
Scatterplot showing credible main effect of bystander age in bonobos (top) and moderate evidence for an interaction between bystander age and sex in chimpanzees (bottom). X-axis = age in years. Y-axis = consolation/total opportunities to console. Females = blue circles; males = red triangles. Point size reflects number of events the individual was present as a bystander. Note, this plot provides a simple visualisation of the aggregated raw data using frequentist 95% confidence limits for ease of comparison. See Figure S3 (ESM) for predicted probability plots.

## Discussion

Systematic cross-species comparisons are vital for elucidating the evolutionary origins for complex socio-emotional behaviours, including empathy. Human research has revealed substantial individual and cultural variation in emotional responding and this study indicates that this may be the case for our closest living relatives. Due to purported increased emotional sensitivity, reduced aggressivity, and higher levels of social tolerance [33,41], it has been suggested that bonobos may be more empathic than chimpanzees. However, our results indicate that the two species do not significantly differ in consolation tendency—considered a behavioural expression of empathy—both in terms of the behaviour’s overall occurrence and the number of consolers that respond per event. Our results reflect recent findings that contrast with these previously held assumed species dichotomisations, such as in aggression [77] and social tolerance [59]. We did, however, find several within-species trends in potential drivers of consolatory tendencies. Expressions of empathy and other emotion-related behaviour are known to vary in wild chimpanzee communities of the same subspecies [24,25], which parallels findings across diverse human cultures [4]. The presence of intraspecies group variation in consolation in the sanctuary bonobos complements this literature and suggests that empathic tendencies may be shaped by broader individual and socio-cultural circumstances.

Our findings complement previous consolation research in *Pan* by showing that bystander age is an important predictor of consolation in bonobo and chimpanzee communities [19,27]. Although underlying mechanisms cannot be detected in an observational study, it is possible that young individuals may console more than older individuals as they are typically afforded more social tolerance before they become fully embedded into their respective group’s dominance dynamics [78]. Furthermore, younger individuals may be less inhibitive with their responses in these contexts. We did not measure individual’s capacity for inhibition in this study. However, an interesting follow-up would be to compare these two species on inhibition, such as through delayed gratification tasks [79], and investigate if this measure predicts an individual’s relative tendency to console.

Consistent with the theory that social closeness predicts empathic responses [2,80], kinship predicted consolation in both species. However, we were only able to test dyadic affiliation levels in chimpanzees, finding that consolation was biased towards closer social partners generally. Previous investigations on these bonobo groups revealed strong influences of social relationship on consolation occurrences [19]. Temporal variation in consolatory efforts may reflect empathy as a phenomenon that emerges flexibly when social conditions permit [13].

It is possible that consolation may serve different functions in bonobo and chimpanzee societies. Chimpanzees may be more selective in their responses as wild and captive communities have been well-documented to exhibit political social lives, centred around strong adult male-male alliances and displays of dominance and subordinacy [33]. This may be further evidenced by a possible age-sex interaction in chimpanzees, whereby young males were the most consolatory and older individuals were the least consolatory. Enhanced responsiveness in young male chimpanzees may represent a drive to form and reinforce strong social bonds with their close peers and older individuals they will spend their lives with [81]. Furthermore, adult male chimpanzees were much less likely to be followed as distressed focals (*N* = 17/160) compared to adult females (*N* = 103/160). Therefore, opportunities for adult male chimpanzees to console their closest social partners were respectively low and may additionally explain the direction of the age-sex interaction whereby older males as unlikely to console as adult females. In the Kanyawara community at Kibale Chimpanzee Project in Uganda, young male chimpanzees are typically more likely to be victims of aggression, which appears to be due to their tendencies to elicit the most aggression themselves [82]. Regardless, social bonds formed during male chimpanzee development predict dominance trajectories [83]. Thus, this effect at a young age in chimpanzees may be caused by younger males watching their peers being aggressed often during a period when they are building foundational relationships for later life [84].

In contrast, in bonobos, males typically only form strong relationships with their mothers [85] and occasionally other females for mating and alliance formation [61]. Despite males being the philopatric sex for both species, only chimpanzee communities tend to feature strong male-male relationships [33]. Furthermore, the political dynamics in bonobo societies appear to be relatively more subtle, and thus difficult to study [54]. Instead, bonobos may be less selective and respond out of care and emotional sensitivity towards others [35,86]. Bonobos tended to console victims of both sexes and all ranks evenly. Furthermore, in line with recent findings that paedomorphic signalling increases likelihood of receiving consolation [87], younger bonobo victims were generally more likely to be consoled. However, higher-ranked bonobo bystanders did offer consolation more than those of low rank, indicating that lower social standing may inhibit individuals to approach distressed peers. A deeper, longitudinal comparison with more groups of each species is needed to reveal the extent to which consolation has a political association across age, rank, and sex combinations in each species.

In this study, we imposed strict criteria of inclusion for consolation, where any behaviour following a solicitation from the victim was not counted. For example, if a victim immediately approached a bystander and solicited an interaction with them, any subsequent interactions initiated by the bystander during the PC/PD period were not coded as consolatory approaches. As consolation is linked with empathy, if bystanders are not the party to solicit contact then it is not possible to infer that they may have been motivated by the victim’s distress to respond [43,88]. Therefore, there is also a possibility that victim-initiated interactions provided reassurance and comfort to the victim, yet these encounters were not counted in our analyses. A wider investigation into bystander-victim interactions following distress could reveal whether the intraspecies trends found in this study are associated with empathic tendencies, or wider conflict management responses in general [89].

Intraspecies variation is vital for comparative research across taxa [90]. Studies of primates have revealed substantial intergroup behavioural variation, such as bartering in long-tailed macaques (*Macaca fascicularis*; [91]), activity and ranging in guerezas (*Colobus guereza*; [92]), and even the connectedness of social networks in vervets (*Chlorocebus pygerythrus*; [93]). The *Pan* apes similarly can vary between groups regarding expressions of various social and ecological behaviours. Examples include, but are by no means limited to, grooming traditions [94] and communication in chimpanzees [95], and hunting behaviour and tool use in bonobos [96]. In addition, bonobos and chimpanzees also show marked intraspecies variation in levels of social tolerance [58,59,97], a factor considered to facilitate empathy [43,44].

Whilst there were no overall species differences between the bonobos and chimpanzees in this sample, individual variation and social relationships appear to drive consolation. The *Social Constraints Hypothesis* [43] posits that empathy is facilitated by social tolerance, and has been suggested to explain why some primates express empathic behaviour and others do not [43,44]. Bonobos have previously been purported to be more socially tolerant than chimpanzees [41,56], however recent analyses indicate that greater variation lies at the group-level in the *Pan* apes [59,77]. Consolation appears to be primarily driven by individual and social characteristics, indicating that group compositional changes may lead to variation in empathic tendencies over time, similar to recent studies on social tolerance [98,99]. For example, groups may score differently on consolation when numbers of juveniles and kin-related pairs fluctuate. In our sample, when controlling for all other factors, there was no credible difference between bonobo groups in consolation tendency. To further elucidate any group-level variation in consolation across the *Pan* apes [43,44], future research should assess tolerance levels and risk of aggression of multiple groups and compare these group-specific factors with relative tendencies to console.

In sum, our findings support the notion that within-species community variation among *Pan* behavioural tendencies may be more significant than between-species differences [33,90,100]. Both bonobos and chimpanzees are highly flexible and adaptable species, and under particular conditions and pressures may exhibit greater or reduced tendencies to express empathy-related behaviours like consolation. In humans, we see individual- and group-level variation in behaviours such as communication, prosociality, conformity, and empathy and other socio-emotional responses that may be facilitated by certain social dynamics and cultural norms [4,101–103]. The same may be true in our closest living relatives. Further research into species comparisons should always integrate group-level differences (e.g., composition, collective temperaments) that may promote or hinder empathic expressions such as consolation. Comparing more groups of the same species and investigating within-group drivers of various empathy-related behaviours may reveal a deep ancestral history of such cultural flexibility in emotional responsiveness, and what features promote or hinder the expression of empathy.

## Supporting information

Electronic Supplementary Material

## Acknowledgements

We thank the keepers, veterinarians, and all other staff of Chimfunshi Wildlife Orphanage Trust and Lola ya Bonobo, including Innocent Mulenga, Felix Chinyama, Thomson Mbilishi, Lameck Musaka, Joseph Kasongo, Thalita Calvi, Fanny Minesi, Raphaël Belais, Stany Mokando, and Jean-Claude Nzumbi. We thank Zoë Goldsborough and Heritier Izansone for their assistance with data collection and Emma Doherty and Georgia Sandars for help with reliability coding. We also thank Edwin van Leeuwen for his support and assistance with this project, particularly with social data collection and analysis. Funding was provided by the Templeton World Charity Foundation, and CW would like to thank the Carper Foundation for additional grant support.

## Data accessibility

All data and R code supporting this article are included as electronic supplementary material.

