## Supplementary material for "Within-species variation eclipses between-species differences in *Pan* consolation": Data and R Code: consolation_r script.R

```
#script for:
# within-species variation eclipses between-species differences in pan consolation
# biorxiv

rm(list=ls())

# save workspace ####
save.image("~/twcf19_pan consolation_workspace.RData")
load("~/twcf19_pan consolation_workspace.RData")

# packages ####

#install.packages("NAME")
require(car)
require(lme4)
require(dplyr)
require(plyr)
require(DHARMa)
require(ggplot2)
require(ggokabeito)
require(ggsignif)
require(brms)
require(tidybayes)
require(shinystan)
require(bayestestR)
require(emmeans)
require(nima)
require(sjPlot)

install.packages("cmdstanr", repos = c("https://mc-stan.org/r-packages/", getOption("repos")))
require(cmdstanr)

# set wd
setwd("~/Dropbox/Projects/TWCF19-Consolation/data and code")

# preparing data ####

# event-level dataframe = one row per event, so we know how many per species featured consolation

events.rs=read.table(file="xevent.txt", header=T, sep="\t", stringsAsFactors = T)

head(xevent)
str(xevent)

table(xevent$victim)
table(xevent$aggressor)

# ape-level dataframe = one row per ape including opps and cons

xapes=read.table(file="xapes.txt", header=T, sep="\t", stringsAsFactors = T)

head(xapes)
str(xapes)

# bonobo.rs = every bystander-victim combo within an event w/ random slope columns

bonobo.rs=read.table(file="xbonobo.txt", header=T, sep="\t", stringsAsFactors = T)

head(bonobo.rs)
str(bonobo.rs)

# chimp.rs = every bystander-victim combo within an event w/ random slope columns

chimp.rs=read.table(file="xchimp.txt", header=T, sep="\t", stringsAsFactors = T)

head(chimp.rs)
str(chimp.rs)

###
###
###
###
###
###

# testing species variability ####

## model 1.1: event-level (binomial) ####

# testing species differences in occurrences of 1+ consolatory approaches

### fit model ####

brm.model1.1 <- brm(consolation_contact~species+z.bystanders+
                      (1+z.bystanders||aggressor)+
                      (1+z.bystanders||victim),
                    data=events.rs, family = bernoulli(), 
                    iter = 10000, warmup = 2000,
                    cores = 10, chains = 4, seed = 123,
                    backend = getOption("cmdstanr"),
                    control = list(adapt_delta = 0.999))

summary(brm.model1.1)

launch_shinystan(brm.model1.1) # to retrieve mean, sd, and n_eff for all variables

# extract 89% credibility intervals
ci_89_1.1 <- round(posterior_summary(brm.model1.1, probs = c(0.055, 0.945)),2)
head(print(ci_89_1.1),7)

# probability of direction results
pd_results_1.1 <- pd(brm.model1.1)
print(pd_results_1.1)

### model diagnostics ####
tail(sort(rstan::summary(brm.model1.1$fit)$summary[,"Rhat"])) # are all r hat < 1.01?
head(sort(rstan::summary(brm.model1.1$fit)$summary[,"n_eff"])) # are all effective samples >1000?
pp_check(brm.model1.1, ndraws=1000) # does posterior distribution reflect real observations of response?
plot(brm.model1.1) # how did the chains behave, any divergent transitions etc.
mcmc_plot(brm.model1.1, type = "areas")

# collinearity
# run a standard linear model without the random effects
lm.full.model1.1=lm(consolation_contact~species+z.bystanders,
                    data=events.rs.model1.1)
vif(lm.full.model1.1)

# species does not predict occurrence of at least one consolatory approach

### plotting effects ####

#### binomial (raw data proportion) ####

table(events.rs$group,events.rs$consolation_contact)

model1.1_plot <- aggregate(events.rs$consolation_contact,
                           by = list(species = c(events.rs$species)),
                           FUN = function(x) c(mean = mean(x), sd = sd(x),
                                               n = length(x)))

# a little manipulation as the previous function returns matrices instead of vectors
model1.1_plot <- do.call(data.frame, model1.1_plot)

levels(model1.1_plot$species) <- c("bonobo", "chimpanzee")

# create theme
My_Theme = theme(
  axis.title.x = element_text(size = 16, face = "bold"),
  axis.text.x = element_text(size = 14, face = "bold"),
  axis.title.y = element_text(size = 16, face = "bold"),
  axis.text.y = element_text(size = 14, face = "bold"),
  legend.title = element_text(size = 16, face = "bold"),
  legend.text = element_text(size = 14, face = "bold"))

# colour palette
library(ggokabeito)
palette.colors(palette = "Okabe-Ito")
##         black        orange       skyblue   bluishgreen        yellow 
##     "#000000"     "#E69F00"     "#56B4E9"     "#009E73"     "#F0E442" 
##          blue    vermillion reddishpurple          gray 
##     "#0072B2"     "#D55E00"     "#CC79A7"     "#999999" 

# create label for x axis
specieslabs <- c("bonobo", "chimpanzee")

# and now compute standard error for each group
# rename the columns just for ease of use
model1.1_plot$se <- model1.1_plot$x.sd / sqrt(model1.1_plot$x.n)
colnames(model1.1_plot) <- c("species", "mean", "sd", "n", "se")

limits1.1 <- aes(ymax = model1.1_plot$mean + model1.1_plot$se,
                 ymin = model1.1_plot$mean - model1.1_plot$se)

label.model1.1_plot <- data.frame(group = c("bonobo", "chimpanzee"),
                                  Value = c(6, 9))

plot_1.1_species <- ggplot(data = model1.1_plot, aes(x = species, y = mean, fill = species)) +
  geom_bar(stat = "identity", position = position_dodge(0.9)) +
  geom_errorbar(limits1.1, position = position_dodge(0.9), width = 0.25) +
  scale_x_discrete(labels = specieslabs) +
  ylim(0,1) +
  theme_minimal() +
  theme(legend.position = "none")+
  scale_fill_manual(values = c("#0072B2", "#D55E00"))+
  labs(x = "species", y = "proportion of events with 1+ consolation", fill = "species") +
  My_Theme
plot_1.1_species


#### binomial (predicted probability) ####

model1.1_ce<-conditional_effects(brm.model1.1, prob = 0.89)
model1.1_ce_plot<-model1.1_ce$species
levels(model1.1_ce_plot$species) <- c("bonobo", "chimpanzee")

plot1.1_prob <- ggplot(model1.1_ce_plot, aes(x=species, y=estimate__, color = species)) + 
  geom_errorbar(aes(ymin=estimate__-se__, ymax=estimate__+se__), width=0.5, size=1) +
  geom_point(size=4, shape=18,) + theme_minimal() + xlab("species") + ylab ("probability of 1+ consolation") +
  theme(strip.text = element_text(size = 14)) +
  theme(legend.position = "none") +
  theme(legend.title = element_text(colour="black", size=14, face="bold")) +
  theme(legend.text = element_text(size=14, face="bold")) +
  theme(axis.text.x = "none") +
  scale_color_manual(values = c("#0072B2", "#D55E00")) +
  theme(axis.title.x=element_blank(),
        axis.text.x=element_blank(),
        axis.ticks.x=element_blank()) +
  scale_y_continuous(limits=c(0,1)) +
  My_Theme
plot1.1_prob

#
#
#
#
#

## model 1.2: event-level (poisson) ####

# do more individuals console per event?

### fit model ####

brm.model1.2 <- brm(consolation_contact_count~species+z.bystanders+
                      (1+z.bystanders||aggressor)+
                      (1+z.bystanders||victim),
                    data=events.rs, family = poisson(link="log"),
                    iter = 10000, warmup = 2000,
                    cores = 10, chains = 4, seed = 123,
                    backend = getOption("cmdstanr"),
                    control = list(adapt_delta = 0.999))

summary(brm.model1.2)

launch_shinystan(brm.model1.2) # to retrieve mean, sd, and n_eff for all variables

# extract 89% credibility intervals
ci_89_1.2 <- round(posterior_summary(brm.model1.2, probs = c(0.055, 0.945)),2)
head(print(ci_89_1.2),7)

# probability of direction results
pd_results_1.2 <- pd(brm.model1.2)
print(pd_results_1.2)

### model diagnostics ####
tail(sort(rstan::summary(brm.model1.2$fit)$summary[,"Rhat"])) # are all r hat < 1.01?
head(sort(rstan::summary(brm.model1.2$fit)$summary[,"n_eff"])) # are all effective samples >1000?
pp_check(brm.model1.2, ndraws=1000) # does posterior distribution reflect real observations of response?
plot(brm.model1.2) # how did the chains behave, any divergent transitions etc.
mcmc_plot(brm.model1.2, type = "areas")


# collinearity
# run a standard linear model without the random effects
lm.full.model1.2=lm(consolation_contact_count~species+z.bystanders,
                    data=events.rs.model1.2)
vif(lm.full.model1.2)

###
###
###
###
###

# testing within-species variability ####

# removing individuals lacking social data or sufficient opportunities due to long-term removal from group

xconsolation <- subset(xconsolation, victim != "dilolo")
xconsolation <- subset(xconsolation, victim != "kodoro")
xconsolation <- subset(xconsolation, victim != "masya")
xconsolation <- subset(xconsolation, victim != "mikey")
xconsolation <- subset(xconsolation, victim != "mumba")
xconsolation <- subset(xconsolation, bystander != "dilolo")
xconsolation <- subset(xconsolation, bystander != "kodoro")
xconsolation <- subset(xconsolation, bystander != "masya")
xconsolation <- subset(xconsolation, bystander != "mikey")
xconsolation <- subset(xconsolation, bystander != "mumba")
xconsolation <- droplevels(xconsolation)

xbonobo <- subset(xconsolation, group != "c2")
xbonobo <- droplevels(xbonobo)
xchimp <- subset(xconsolation, group == "c2")
xchimp <- droplevels(xchimp)

table(xbonobo$group,xbonobo$consolation_contact)
table(xchimp$consolation_contact)

## model 2.1: bernoulli testing dyad-level within event [bonobo] ####

### fit model ####

# with random slopes
brm.model2.1=brm(consolation_contact ~ group +
                   z.bystander_age * bystander_sex + bystander_rank +
                   z.victim_age + victim_rank + victim_sex +
                   kinship +
                   (1 | aggressor) +
                   (1 + z.victim_age + victim_sex.M || bystander)+
                   (1 | event_id) +
                   (1 + z.bystander_age + bystander_sex.M || victim),
                 data=bonobo.rs, family = bernoulli(),
                 iter = 10000, warmup = 2000,
                 cores = 10, chains = 4, seed = 123,
                 backend = getOption("cmdstanr"),
                 control = list(adapt_delta = 0.999))

summary(brm.model2.1)

launch_shinystan(brm.model2.1) # to retrieve mean, sd, and n_eff for all variables

# probability of direction results
pd_results_2.1 <- pd(brm.model2.1)
print(pd_results_2.1)

# extract 89% credibility intervals
ci_89_2.1 <- round(posterior_summary(brm.model2.1, probs = c(0.055, 0.945)),2)
head(print(ci_89_2.1),20)

### model diagnostics ####
tail(sort(rstan::summary(brm.model2.1$fit)$summary[,"Rhat"])) # are all r hat < 1.01?
head(sort(rstan::summary(brm.model2.1$fit)$summary[,"n_eff"])) # are all effective samples >1000?
pp_check(brm.model2.1, ndraws=1000) # does posterior distribution reflect real observations of response?
plot(brm.model2.1) # how did the chains behave, any divergent transitions etc.
mcmc_plot(brm.model2.1, type = "areas")

# collinearity
# run a standard linear model without the random effects
lm.full.model2.1=lm(consolation_contact ~ group + z.bystander_age + bystander_sex + bystander_rank + 
                      z.victim_age + victim_rank + victim_sex + kinship,
                    data=bonobo.rs)
vif(lm.full.model2.1)

#
#
#
#
#

## model 2.2: bernoulli testing dyad-level within event [chimp] ####

### fit model ####

# with random slopes
brm.model2.2=brm(consolation_contact ~ z.bystander_age * bystander_sex + bystander_rank +
                   z.victim_age + victim_rank + victim_sex +
                   z.dyad_aff + kinship + z.socproxrate +
                   (1 | aggressor) +
                   (1 + z.victim_age + victim_sex.M || bystander)+
                   (1 | event_id) +
                   (1 + z.bystander_age + bystander_sex.M || victim),
                 data=chimp.rs, family = bernoulli(),
                 iter = 10000, warmup = 2000,
                 cores = 10, chains = 4, seed = 123,
                 backend = getOption("cmdstanr"),
                 control = list(adapt_delta = 0.999))

summary(brm.model2.2)

launch_shinystan(brm.model2.2) # to retrieve mean, sd, and n_eff for all variables

# probability of direction results
pd_results_2.2 <- pd(brm.model2.2)
print(pd_results_2.2)

# extract 89% credibility intervals
ci_89_2.2 <- round(posterior_summary(brm.model2.2, probs = c(0.055, 0.945)),2)
head(print(ci_89_2.2),21)

### model diagnostics ####
tail(sort(rstan::summary(brm.model2.2$fit)$summary[,"Rhat"])) # are all r hat < 1.01?
head(sort(rstan::summary(brm.model2.2$fit)$summary[,"n_eff"])) # are all effective samples >1000?
pp_check(brm.model2.2, ndraws=1000) # does posterior distribution reflect real observations of response?
plot(brm.model2.2) # how did the chains behave, any divergent transitions etc.
mcmc_plot(brm.model2.2, type = "areas")

# collinearity
# run a standard linear model without the random effects
lm.full.model2.2=lm(consolation_contact ~ z.bystander_age + bystander_sex + bystander_rank + 
                      z.victim_age + victim_rank + victim_sex + z.dyad_aff + kinship + z.socproxrate,
                    data=chimp.rs)
vif(lm.full.model2.2)

#
#
#
#
#

## plotting ####

### kinship ####

kinship_b<-conditional_effects(brm.model2.1, prob = 0.89, effects = "kinship")
kinship_c<-conditional_effects(brm.model2.2, prob = 0.89, effects = "kinship")
kinship_b_plot<-kinship_b$kinship
kinship_c_plot<-kinship_c$kinship
levels(kinship_b_plot$kinship) <- c("kin", "nonkin")
levels(kinship_c_plot$kinship) <- c("kin", "nonkin")

My_Theme = theme(
  title = element_text(size = 18, face = "bold"),
  axis.title.y = element_text(size = 18, face = "bold"),
  axis.text.y = element_text(size = 16),
  legend.title = element_text(size = 18, face = "bold"),
  legend.text = element_text(size = 16))

plot_kin_b <- ggplot(kinship_b_plot, aes(x=kinship, y=estimate__, color = kinship)) + 
  geom_errorbar(aes(ymin=estimate__-se__, ymax=estimate__+se__), width=0.5, size=1) +
  geom_point(size=4, shape=18,) + theme_minimal() + xlab("kinship") + ylab ("predicted probability of consolation") +
  theme(strip.text = element_text(size = 14)) +
  theme(legend.position = "bottom") +
  theme(legend.title = element_text(colour="black", size=14, face="bold")) +
  theme(legend.text = element_text(size=14, face="bold")) +
  theme(axis.text.x = "none") +
  scale_color_manual(values = c("#0072B2", "#D55E00")) +
  theme(axis.title.x=element_blank(),
        axis.text.x=element_blank(),
        axis.ticks.x=element_blank()) +
  scale_y_continuous(limits=c(0,NA)) +
  ggtitle("(1a) bonobos") +
  My_Theme
plot_kin_b

plot_kin_c <- ggplot(kinship_c_plot, aes(x=kinship, y=estimate__, color = kinship)) + 
  geom_errorbar(aes(ymin=estimate__-se__, ymax=estimate__+se__), width=0.5, size=1) +
  geom_point(size=4, shape=18,) + theme_minimal() + xlab("kinship") + ylab ("predicted probability of consolation") +
  theme(strip.text = element_text(size = 14)) +
  theme(legend.position = "bottom") +
  theme(legend.title = element_text(colour="black", size=14, face="bold")) +
  theme(legend.text = element_text(size=14, face="bold")) +
  theme(axis.text.x = "none") +
  scale_color_manual(values = c("#0072B2", "#D55E00")) +
  theme(axis.title.x=element_blank(),
        axis.text.x=element_blank(),
        axis.ticks.x=element_blank()) +
  scale_y_continuous(limits=c(0,NA)) +
  ggtitle("(1b) chimpanzees") +
  My_Theme
plot_kin_c

#
#
#
#
#

### dyadic aff ####

# generate conditional effects
conditions_c <- conditional_effects(brm.model2.2, effects = "z.dyad_aff", 
                                    prob = 0.89)

# extract the data
plot_data_c <- as.data.frame(conditions_c$`z.dyad_aff`)

My_Theme = theme(
  title = element_text(size = 20, face = "bold"),
  axis.title.x = element_text(size = 20, face = "bold"),
  axis.text.x = element_text(size = 18),
  axis.title.y = element_text(size = 20, face = "bold"),
  axis.text.y = element_text(size = 18),
  legend.title = element_text(colour="black", face="bold"),
  legend.text = element_text(size = 18),
  legend.position = "bottom")

# plot
plot_dyad_aff_c <- ggplot(plot_data_c, aes(x = z.dyad_aff, y = estimate__)) +
  geom_line() +
  geom_ribbon(aes(ymin = lower__, ymax = upper__), alpha = 0.5) +
  theme_nima() +
  theme(legend.position="bottom") +
  labs(x = "dyadic affiliation level", y = "predicted probability of consolation") +
  ggtitle("(2) chimpanzees") +
  theme_minimal() +
  theme(legend.position = "top") +
  My_Theme
plot_dyad_aff_c

#
#
#
#
#

### victim age ####

# generate conditional effects
conditions_b <- conditional_effects(brm.model2.1, effects = "z.victim_age", 
                                    prob = 0.89)
conditions_c <- conditional_effects(brm.model2.2, effects = "z.victim_age", 
                                    prob = 0.89)

# extract the data
plot_data_b <- as.data.frame(conditions_b$`z.victim_age`)
plot_data_c <- as.data.frame(conditions_c$`z.victim_age`)

My_Theme = theme(
  title = element_text(size = 20, hjust = 0.5, face = "bold"),
  axis.title.x = element_text(size = 20, face = "bold"),
  axis.text.x = element_text(size = 18),
  axis.title.y = element_text(size = 20, face = "bold"),
  axis.text.y = element_text(size = 18),
  legend.title = element_text(colour="black", face="bold"),
  legend.text = element_text(size = 18),
  legend.position = "bottom")

# plot
plot_vage_b <- ggplot(plot_data_b, aes(x = z.victim_age, y = estimate__)) +
  geom_line() +
  geom_ribbon(aes(ymin = lower__, ymax = upper__), alpha = 0.5) +
  theme_nima() +
  theme(legend.position="bottom") +
  labs(x = "victim age", y = "predicted probability of consolation") +
  ylim(0,NA) +
  ggtitle("bonobos") +
  theme_minimal() +
  theme(legend.position = "top") +
  My_Theme
plot_vage_b

plot_vage_c <- ggplot(plot_data_c, aes(x = z.victim_age, y = estimate__)) +
  geom_line() +
  geom_ribbon(aes(ymin = lower__, ymax = upper__), alpha = 0.5) +
  theme_nima() +
  theme(legend.position="bottom") +
  labs(x = "victim age", y = "predicted probability of consolation") +
  ylim(0,NA) +
  ggtitle("chimpanzees") +
  theme_minimal() +
  theme(legend.position = "top") +
  My_Theme
plot_vage_c

#
#
#
#
#

### interaction ####

#### raw data ####

# create plot y value
xapes$consrate<-xapes$consolations/xapes$cons_opportunities
xapes_plot<-subset(xapes, cons_opportunities != 0)

bonobo_plot<-subset(xapes_plot, species == "bonobo")
bonobo_plot<-subset(bonobo_plot, bystander != "dilolo")
bonobo_plot<-subset(bonobo_plot, bystander != "kodoro")
bonobo_plot<-droplevels(bonobo_plot)

chimp_plot<-subset(xapes_plot, species == "chimpanzee")
chimp_plot<-subset(chimp_plot, bystander != "masya")
chimp_plot<-subset(chimp_plot, bystander != "mikey")
chimp_plot<-subset(chimp_plot, bystander != "mumba")
chimp_plot<-droplevels(chimp_plot)

My_Theme = theme(
  title = element_text(size = 20, hjust = 0.5, face = "bold"),
  axis.title.x = element_text(size = 20, face = "bold"),
  axis.text.x = element_text(size = 18),
  axis.title.y = element_text(size = 20, face = "bold"),
  axis.text.y = element_text(size = 18),
  legend.title = element_text(colour="black", face="bold"),
  legend.text = element_text(size = 18),
  legend.position = "bottom")

##### bonobos ####
require(ggplot2)
model3_plot_int <- ggplot(data = bonobo_plot, aes(x = age, y = consrate, fill = sex, shape = sex)) +
  geom_point(data = bonobo_plot, mapping = aes(size = cons_opportunities, color = sex), alpha = .5) +
  stat_smooth(data = bonobo_plot, method = "lm", mapping = aes(color = sex), se = TRUE, size = 1, alpha = .5, level = 0.95) +
  scale_fill_manual(values = c("#0072B2", "#D55E00")) +
  scale_colour_manual(values = c("#0072B2", "#D55E00")) +
  theme_nima() +
  ggtitle("bonobos") +
  labs(x = "bystander age", y = "consolatory tendency", 
       fill = "bystander sex", shape = "bystander sex", 
       color = "bystander sex", size = "opportunities") +
  ylim(-0.05,0.2) +
  xlim(0,45) +
  My_Theme
model3_plot_int

##### chimps ####
require(nima)
model4_plot_int <- ggplot(data = chimp_plot, aes(x = age, y = consrate, fill = sex, shape = sex)) +
  geom_point(data = chimp_plot, mapping = aes(size = cons_opportunities, color = sex), alpha = .5) +
  stat_smooth(data = chimp_plot, method = "lm", mapping = aes(color = sex), se = TRUE, size = 1, alpha = .5, level = 0.95) +
  scale_fill_manual(values = c("#0072B2", "#D55E00")) +
  scale_colour_manual(values = c("#0072B2", "#D55E00")) +
  theme_nima() +
  theme(legend.position="bottom") +
  theme(legend.title = element_text(colour="black", face="bold"))+
  theme(axis.title.x = element_text(face = "bold")) +
  theme(axis.title.y = element_text(face = "bold")) +
  theme(plot.title = element_text(hjust = 0.5, face = "bold")) +
  ggtitle("chimpanzees") +
  labs(x = "bystander age", y = "consolatory tendency", 
       fill = "bystander sex", shape = "bystander sex", 
       color = "bystander sex", size = "opportunities") +
  scale_y_continuous(breaks = seq(-0.05,0.2, by = 0.025)) +
  scale_x_continuous(breaks = seq(0,45, by = 5)) +
  ylim(-0.05,0.2) +
  xlim(0,45) +
  guides(size = guide_legend(nrow = 2)) +
  My_Theme
model4_plot_int

#
#
#
#
#

#### conditional plots ####

# generate conditional effects
conditions_b <- conditional_effects(brm.model2.1, effects = "z.bystander_age:bystander_sex", 
                                    prob = 0.89)
conditions_c <- conditional_effects(brm.model2.2, effects = "z.bystander_age:bystander_sex", 
                                    prob = 0.89)

# extract the data
plot_data_b <- as.data.frame(conditions_b$`z.bystander_age:bystander_sex`)
plot_data_c <- as.data.frame(conditions_c$`z.bystander_age:bystander_sex`)

# plot
##### bonobos ####
interaction_plot_b <- ggplot(plot_data_b, aes(x = z.bystander_age, y = estimate__, 
                                              color = bystander_sex, fill = bystander_sex)) +
  geom_line() +
  geom_ribbon(aes(ymin = lower__, ymax = upper__), alpha = 0.5) +
  scale_fill_manual(values = c("#0072B2", "#D55E00")) +
  scale_colour_manual(values = c("#0072B2", "#D55E00")) +
  theme_nima() +
  theme(legend.position="bottom") +
  labs(x = "bystander age", y = "predicted probability of consolation", 
       fill = "bystander sex",
       color = "bystander sex") +
  theme_minimal() +
  theme(legend.position = "top") +
  ggtitle("bonobos")
interaction_plot_b

##### chimpanzees ####
interaction_plot_c <- ggplot(plot_data_c, aes(x = z.bystander_age, y = estimate__, 
                                              color = bystander_sex, fill = bystander_sex)) +
  geom_line() +
  geom_ribbon(aes(ymin = lower__, ymax = upper__), alpha = 0.5) +
  scale_fill_manual(values = c("#0072B2", "#D55E00")) +
  scale_colour_manual(values = c("#0072B2", "#D55E00")) +
  theme_nima() +
  theme(legend.position="bottom") +
  labs(x = "bystander age", y = "predicted probability of consolation", 
       fill = "bystander sex",
       color = "bystander sex") +
  theme_minimal() +
  theme(legend.position = "top") +
  ggtitle("chimpanzees")
interaction_plot_c
```
