## Supplementary material for "Within-species variation eclipses between-species differences in *Pan* consolation": Electronic Supplementary Material

**Table S1. Social compositions^a^ of studied groups.** Bonobos observed at Lola ya Bonobo Sanctuary in the Democratic Republic of the Congo and chimpanzees observed at Chimfunshi Wildlife Orphanage Trust in Zambia.

| **Species** | **Group** | **Total** | **Age Range (in years)** | **Sex** |
| --- | --- | --- | --- | --- |
|  |  |  |  | **F / M^b^** |
| Bonobos  *(Pan paniscus)* | 1 (B1) | 22 | 2–24 | 12 / 10 |
|  | 2 (B2) | 18 | 3–30 | 8 / 10 |
| Chimpanzees  *(Pan troglodytes)* | 2 (C2) | 50 | 1–42 | 33 / 17 |
| Total |  | 90 | 1–42 | 53 / 37 |
| *^a^ Table includes all individuals belonging to their respective groups from the start of observations, all of whom were considered for the main species comparison models. For the within-species models (Models 2.1–2.2), N=2 bonobos and N=3 chimpanzees were removed due to limited observations.*  *^b^ F = Number of females; M = Number of males* | | | | |

**Table S2. Ethogram of affiliative contact behaviours used during post-conflict interactions.** [1–4]

| Behaviour | Description |
| --- | --- |
| **Body kiss | Kiss or soft gnaw on the recipient’s body or burying face in other’s fur. Not within a play context. |
| Contact sit | Sitting in physical contact with another individual. Not engaged in any other social behaviour. |
| Embrace | Gently placing one arm around the other’s shoulder, back or waist, or putting both arms around the other. May include pulling him or her closer. Includes lateral, ventral and side hugging/embracing. |
| **Finger/hand in mouth | Placement of one’s finger or hand in another individual’s mouth (actor is one who extends hand/finger). |
| Genital inspection | Individual touches genitals of another with hand or foot. May sniff finger or sniff genitals directly but only coded if contact occurs. Sniffing not coded without contact. May groom. May shake penis. Includes poking. |
| *Genito-genital contact^a^ | Ventro-ventral embrace. Individuals swing their hips laterally, while keeping vulvae in contact. Not common in chimpanzees. |
| Grasp hand | Individual grasps hand of another individual, usually without shaking motion. |
| Groom | Pushing the hair of another individual back with the thumb or index finger of one hand and holding it back while picking at the exposed skin with the nail of the thumb, index finger, teeth, or lips of the other. ≥ 5sec duration; score only first interaction per grooming session; new event only after 30sec w/o grooming. |
| Hunch-over | One arm or the entire upper body is moved over or briefly pressed on the crouching partner. Not hugging/embracing. |
| Mount | Dorsal-ventral embrace. Individual touches or holds another’s back or sides with or without intromission. May include thrusting pelvis movements. May include body kissing. |
| Mouth kiss | Kissing in mouth-nose region. |
| *Mount walk^b^ | Resting head, arms or chest on the back of another individual while moving forward. |
| Pat^c^ | Repeatedly tapping a hand on another individual’s body. |
| Play | Any instances of playful behaviour with a partner (including wrestling and tickling). May or may not be accompanied by play faces and/or laughter. |
| **Rump-rump touch^a^ | Individuals face away from each other while putting their rumps in contact. Usually, no swinging or rubbing. |
| Touch | Other gentle hand or foot contact, including holding. Laying palm or fingers on another individual. |
| ** Not observed during chimpanzee events*  *** Not observed during bonobo events*  *^a^ “Genito-genital contact” and “Rump-rump touch” merged due to similarity in form*  *^b^ Coded separately but merged with “Mount” due to rarity of occurrences*  *^c^ Coded separately but merged with “Touch” due to rarity of occurrences* | |

**Table S3**. Summary of Bayesian generalised linear mixed models testing species differences in consolation tendencies. Dependent variables are listed in bold font in header lines. Abbreviation: *eff. N* = Effective Sample Size; *b* = estimate; *S.D.* = standard deviation.

| **Model 1.1 Consolation occurrence (observations *N* = 276), Bernoulli distribution (did consolation occur 1/0)** | | | | | |
| --- | --- | --- | --- | --- | --- |
|  | *eff. N* | *b* | *S.D.* | *5.50%* | *94.50%* |
| Intercept | 20,943 | -0.7 | 0.25 | -1.07 | 0.45 |
| Species [chimpanzee]^a^ | 22,849 | 0.08 | 0.33 | -0.94 | 1.11 |
| No. bystanders^b^ | 20,654 | 0.13 | 0.15 | -0.36 | 0.62 |
| Aggressor Intercept | 11,746 | 0.38 | 0.14 | 0.45 | 1.26 |
| Victim Intercept | 5,795 | 0.41 | 0.16 | 0.26 | 1.46 |
| Slope [No. bystanders \| Aggressor] | 10,542 | 0.18 | 0.13 | 0.03 | 0.79 |
| Slope [No. bystanders \| Victim] | 5,663 | 0.26 | 0.15 | 0.07 | 1.21 |
| **Model 1.2 Consolatory approach count (observations *N* = 276), Poisson distribution (number of bystander-initiated interactions as counts)** | | | | | |
|  | *eff. N* | *b* | *S.D.* | *5.50%* | *94.50%* |
| Intercept | 20,943 | -0.7 | 0.25 | -1.12 | -0.30 |
| Species [chimpanzee]^a^ | 22,849 | 0.08 | 0.33 | -0.44 | 0.60 |
| No. bystanders^b^ | 20,654 | 0.13 | 0.15 | -0.12 | 0.37 |
| Aggressor Intercept | 11,746 | 0.38 | 0.14 | 0.18 | 0.61 |
| Victim Intercept | 5,795 | 0.41 | 0.16 | 0.13 | 0.65 |
| Slope [No. bystanders \| Aggressor] | 10,542 | 0.18 | 0.13 | 0.02 | 0.42 |
| Slope [No. bystanders \| Victim] | 5,663 | 0.26 | 0.15 | 0.03 | 0.51 |
| *^a^ reference level = bonobo*  *^b^ z-transformed to a mean of 0 and a standard deviation of 1* | | | | | |

**Table S4**. Summary of Bayesian generalised linear mixed models testing within-species differences in consolation tendencies. Dependent variables are listed in bold font in header lines. Abbreviation: *eff. N* = Effective Sample Size; *b* = estimate; *S.D.* = standard deviation.

| **Model 2.1 Bonobo model (observations *N* = 1656), Bernoulli distribution (did consolation occur 1/0)** | | | | | |
| --- | --- | --- | --- | --- | --- |
|  | *eff. N* | *b* | *S.D.* | *5.50%* | *94.50%* |
| Intercept | 32,000 | -1.68 | 0.71 | -2.84 | -0.57 |
| Group [B2] | 32,000 | -0.49 | 0.52 | -1.34 | 0.32 |
| Bystander age | 32,000 | -1.02 | 0.31 | -1.52 | -0.53 |
| Bystander sex [male] | 32,000 | -0.55 | 0.52 | -1.40 | 0.25 |
| Bystander age * bystander sex [male] | 32,000 | -0.25 | 0.53 | -1.12 | 0.56 |
| Bystander rank [low] | 32,000 | -1.01 | 0.50 | -1.81 | -0.24 |
| Bystander rank [medium] | 32,000 | -0.43 | 0.51 | -1.24 | 0.37 |
| Victim age | 19,564 | -0.56 | 0.29 | -1.06 | -0.13 |
| Victim rank [low] | 32,000 | 0.11 | 0.49 | -0.65 | 0.89 |
| Victim rank [medium] | 32,000 | 0.15 | 0.56 | -0.75 | 1.05 |
| Victim sex [male] | 32,000 | 0.01 | 0.51 | -0.80 | 0.83 |
| Kinship [non-kin] | 32,000 | -2.35 | 0.62 | -3.37 | -1.37 |
| Aggressor Intercept | 11,977 | 0.77 | 0.40 | 0.21 | 1.47 |
| Bystander Intercept | 9,875 | 0.39 | 0.28 | 0.04 | 0.89 |
| Event ID Intercept | 12,156 | 0.32 | 0.23 | 0.03 | 0.73 |
| Victim Intercept | 15,733 | 0.28 | 0.22 | 0.02 | 0.68 |
| Slope [Victim age \| Bystander] | 8,800 | 0.59 | 0.29 | 0.13 | 1.07 |
| Slope [Victim sex \| Bystander] | 10,311 | 1.52 | 0.55 | 0.67 | 2.42 |
| Slope [Bystander age \| Victim] | 16,370 | 0.26 | 0.20 | 0.02 | 0.63 |
| Slope [Bystander sex \| Victim] | 15,135 | 0.59 | 0.43 | 0.05 | 1.38 |
| **Model 2.2 Chimpanzee model (observations *N* = 5668), Bernoulli distribution (did consolation occur 1/0)** | | | | | |
|  | *eff. N* | *b* | *S.D.* | *5.50%* | *94.50%* |
| Intercept | 21,298 | -4.27 | 0.81 | -5.58 | -3.02 |
| Bystander age | 20,938 | -0.67 | 0.29 | -1.14 | -0.23 |
| Bystander sex [male] | 21,430 | 0.08 | 0.62 | -0.90 | 1.05 |
| Bystander age * bystander sex [male] | 23,155 | -0.76 | 0.61 | -1.77 | 0.19 |
| Bystander gregariousness | 23,507 | -0.16 | 0.21 | -0.49 | 0.17 |
| Bystander rank [low] | 19,847 | -0.52 | 0.52 | -1.36 | 0.30 |
| Bystander rank [medium] | 21,602 | -0.50 | 0.51 | -1.31 | 0.32 |
| Victim age | 18,263 | 0.28 | 0.25 | -0.13 | 0.68 |
| Victim rank [low] | 22,423 | 0.21 | 0.57 | -0.71 | 1.11 |
| Victim rank [medium] | 22,295 | 0.43 | 0.61 | -0.53 | 1.41 |
| Victim sex [male] | 20,526 | -0.56 | 0.65 | -1.62 | 0.41 |
| Dyadic affiliation level | 32,000 | 0.20 | 0.09 | 0.05 | 0.35 |
| Kinship [non-kin] | 32,000 | -0.86 | 0.41 | -1.50 | -0.19 |
| Aggressor Intercept | 9,987 | 0.34 | 0.20 | 0.05 | 0.70 |
| Bystander Intercept | 12,020 | 0.93 | 0.24 | 0.59 | 1.34 |
| Event ID Intercept | 5,214 | 0.62 | 0.26 | 0.17 | 1.01 |
| Victim Intercept | 6,926 | 0.67 | 0.31 | 0.17 | 1.19 |
| Slope [Victim age \| Bystander] | 13,683 | 0.18 | 0.13 | 0.02 | 0.42 |
| Slope [Victim sex \| Bystander] | 13,217 | 0.57 | 0.42 | 0.05 | 1.33 |
| Slope [Bystander age \| Victim] | 7,631 | 0.49 | 0.26 | 0.09 | 0.93 |
| Slope [Bystander sex \| Victim] | 8,734 | 1.08 | 0.44 | 0.42 | 1.82 |
| *^a^ z-transformed to a mean of 0 and a standard deviation of 1*  *^b^ reference level = B1*  *^c^ reference level = female*  *^d^ reference level = high*  *^e^ reference level = kin* | | | | | |

**Fig S1** Trace plots of MCMC chains and posterior distributions of all models.

Model 1.1

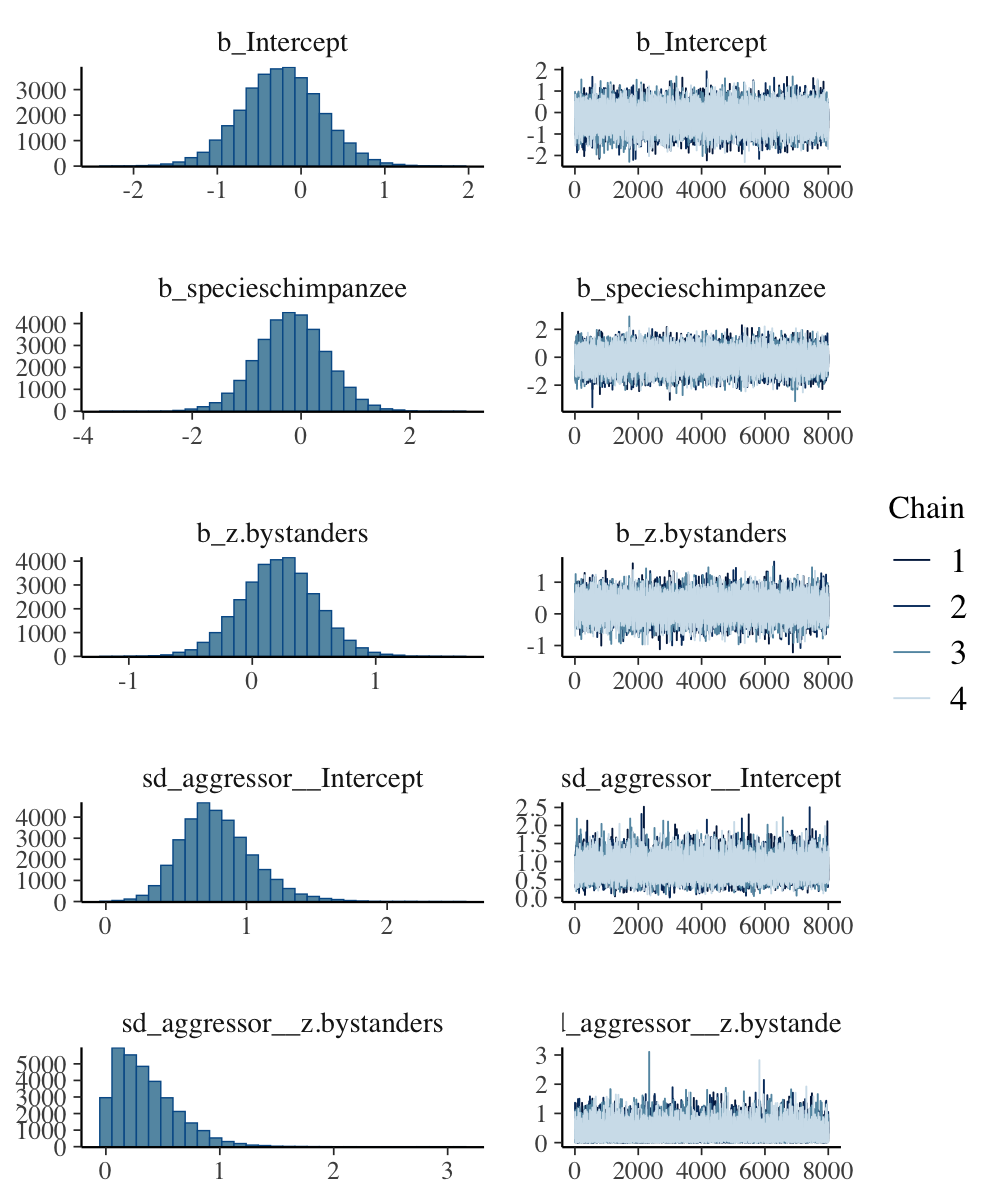

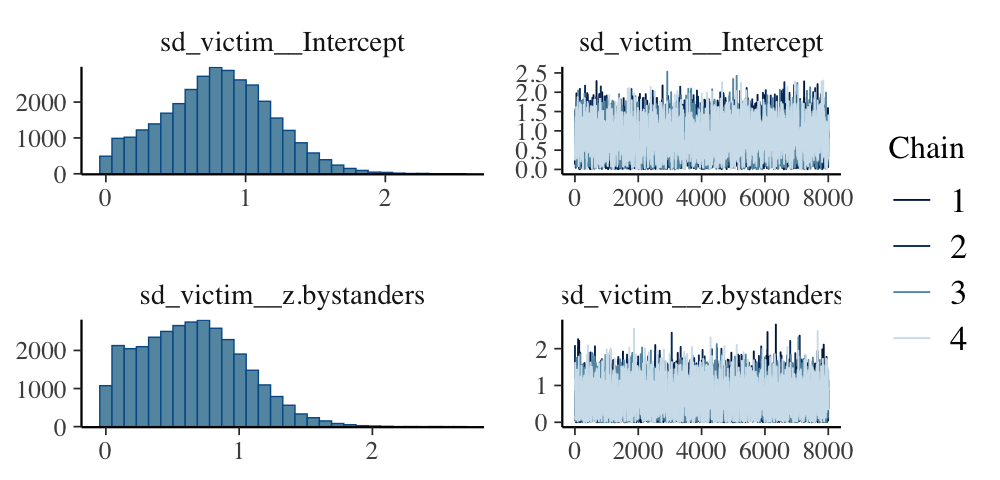

Model 1.2

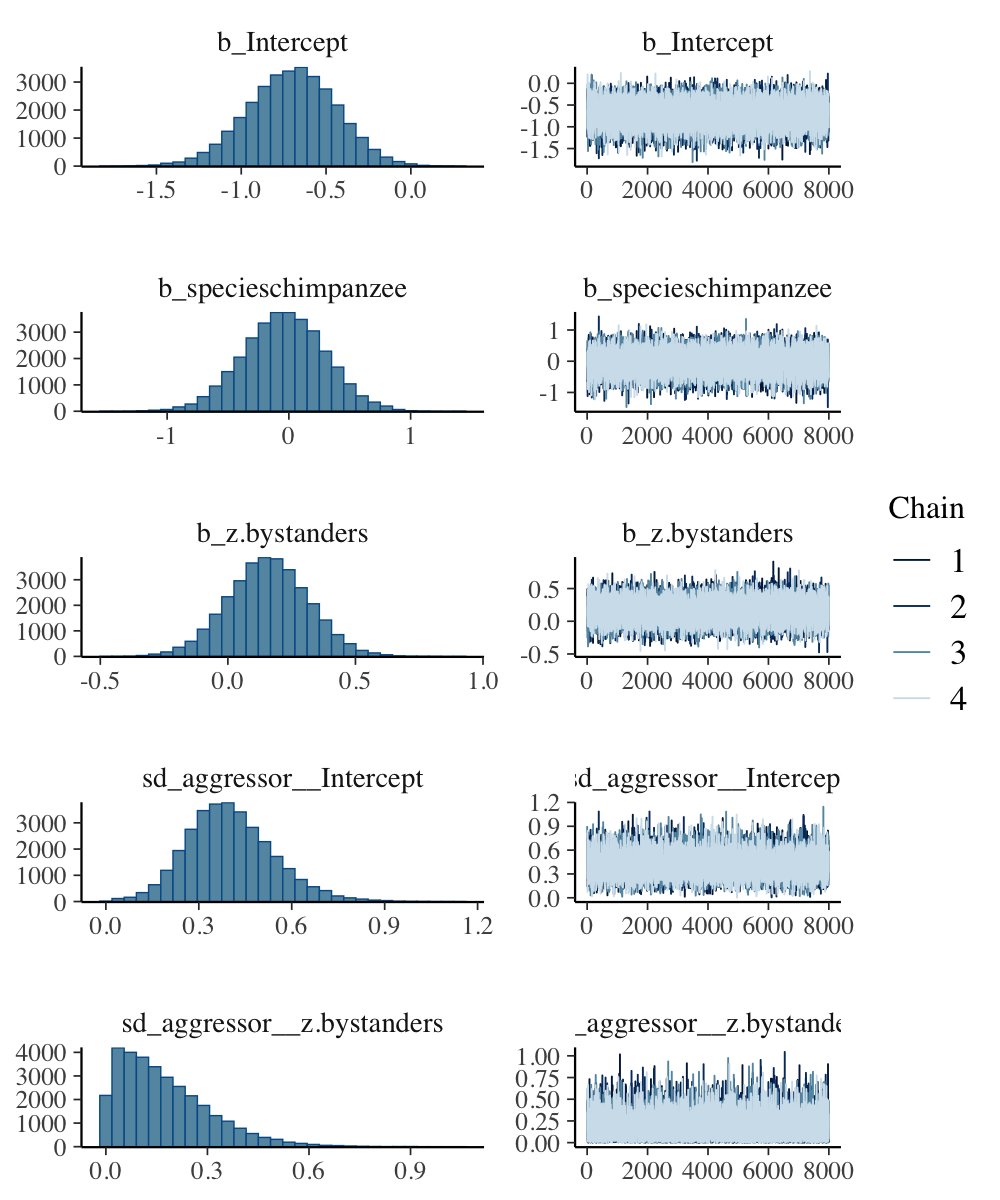

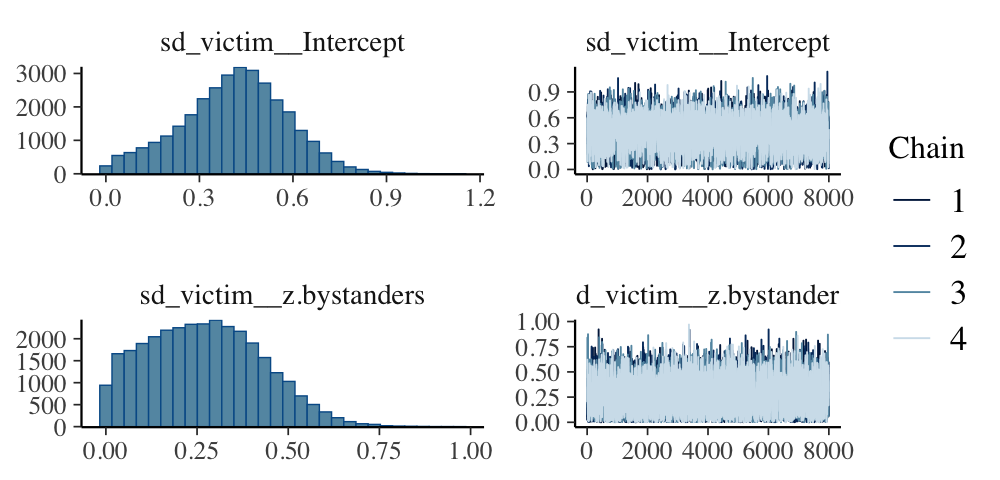

Model 2.1

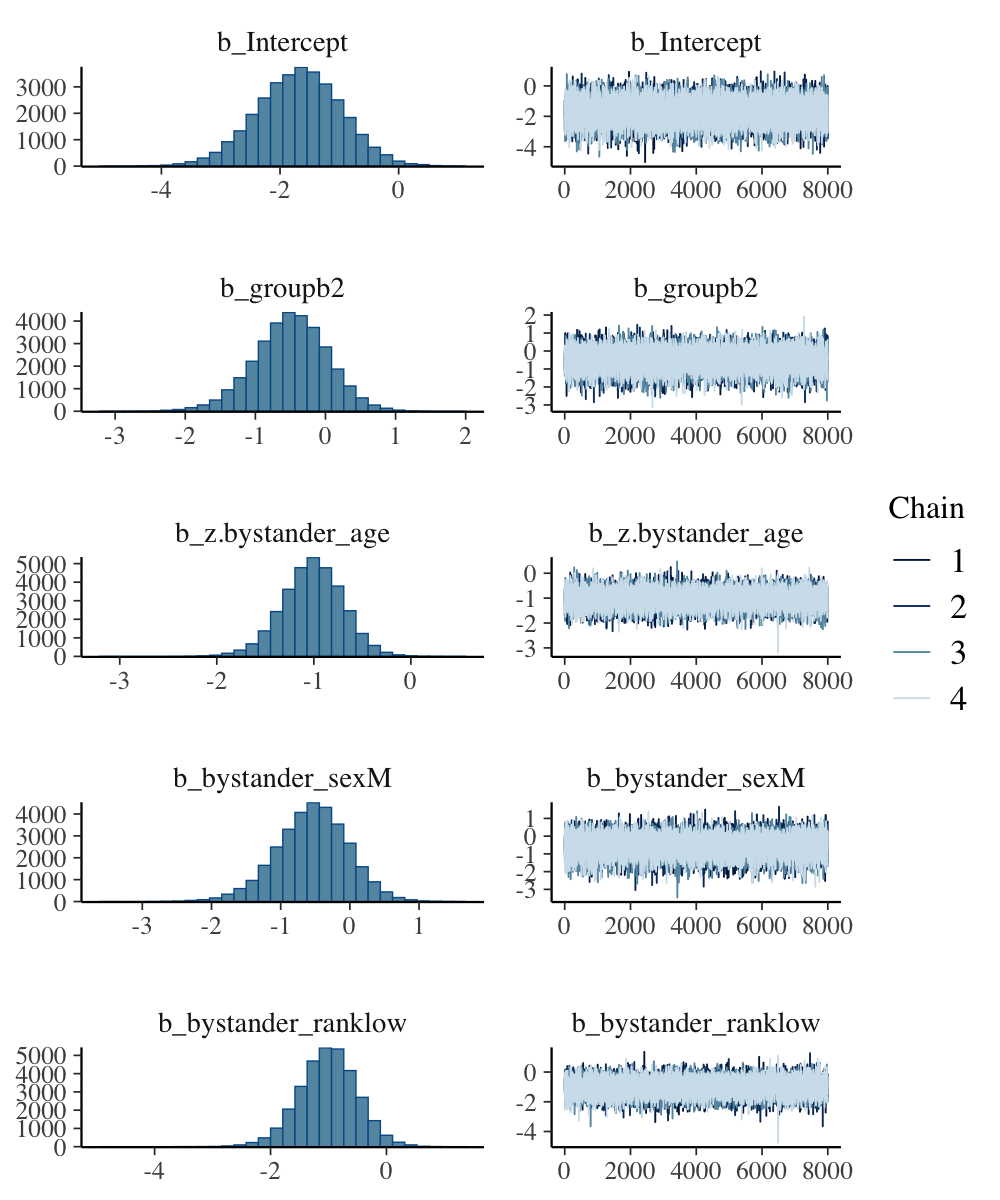

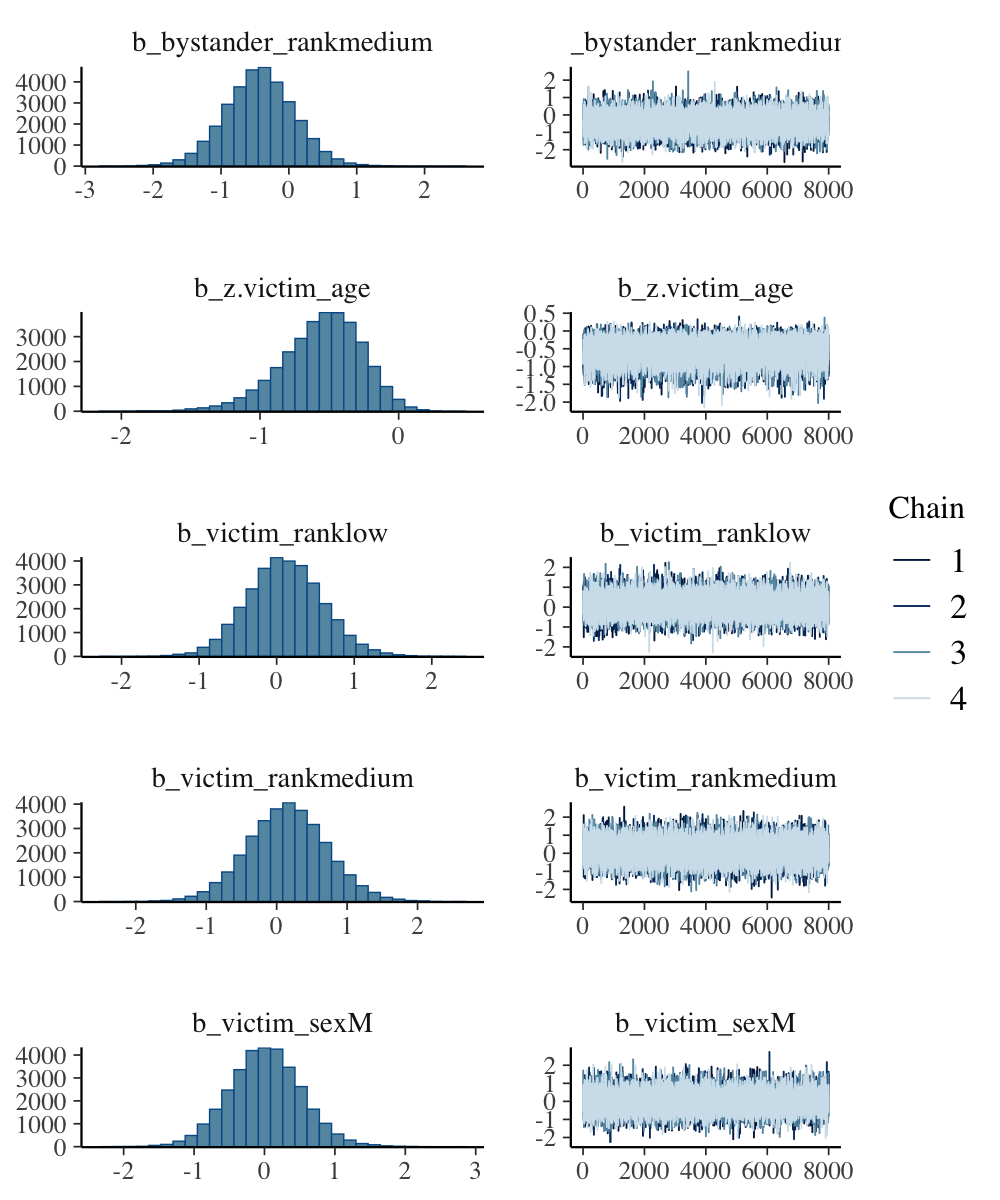

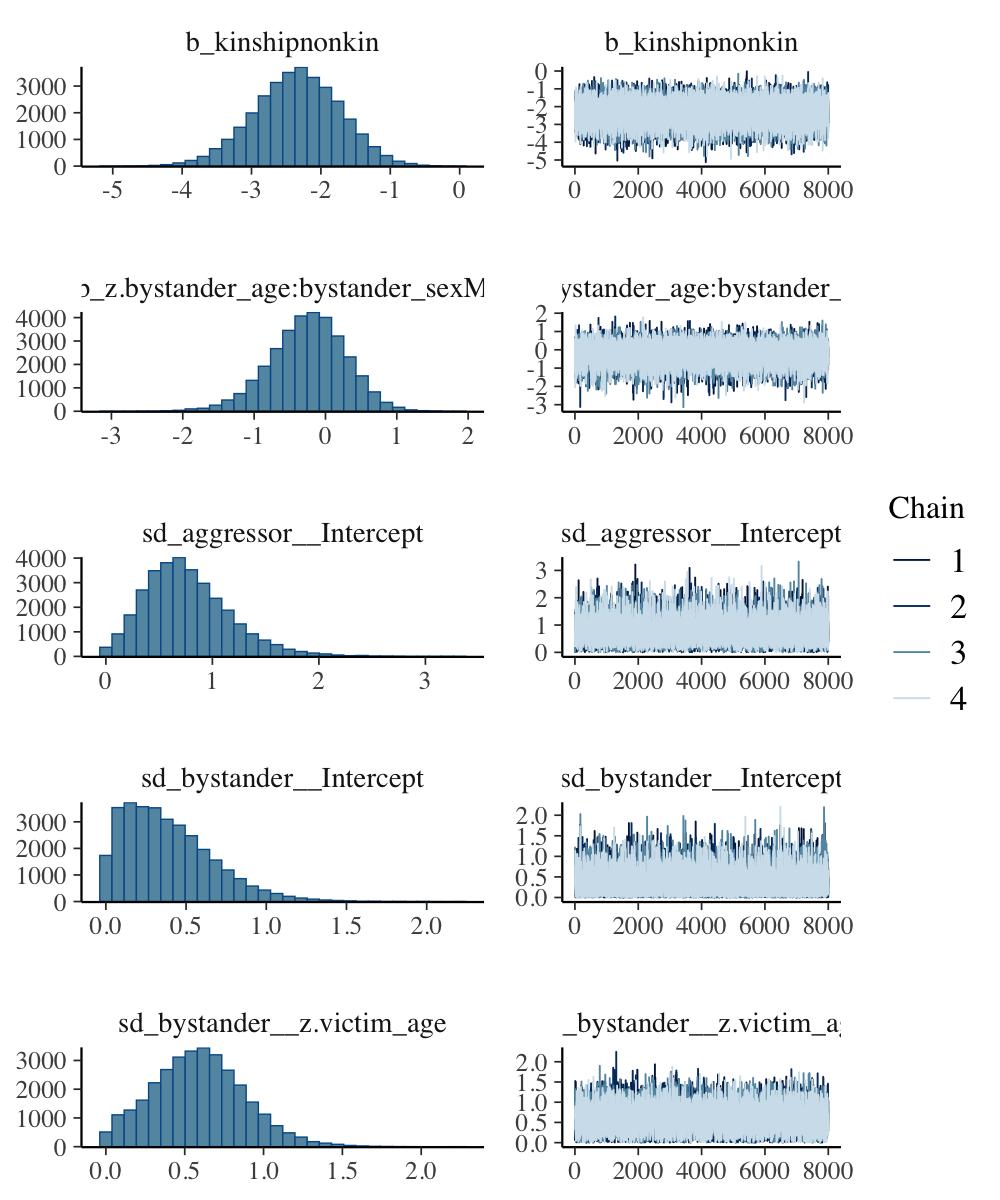

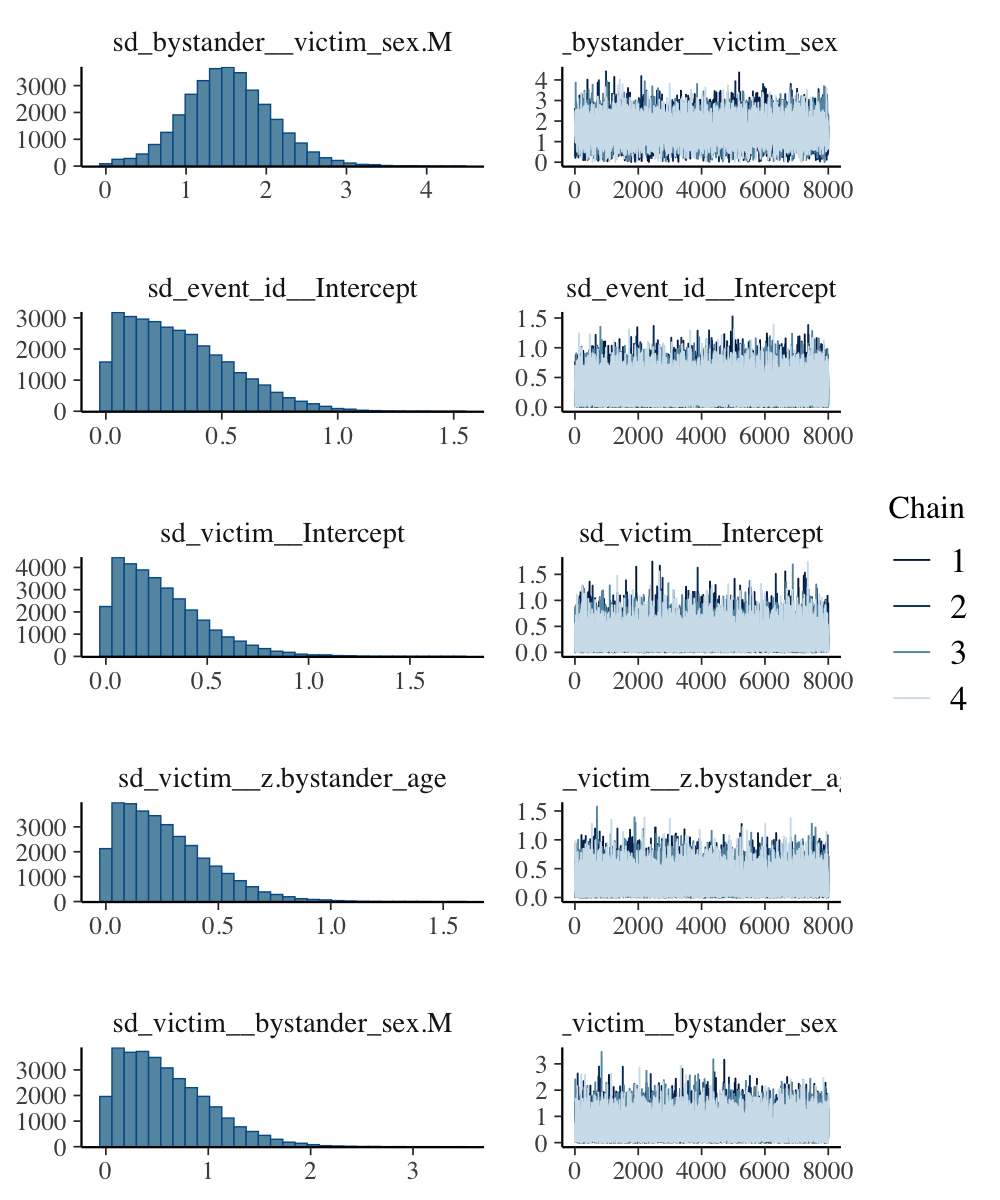

Model 2.2

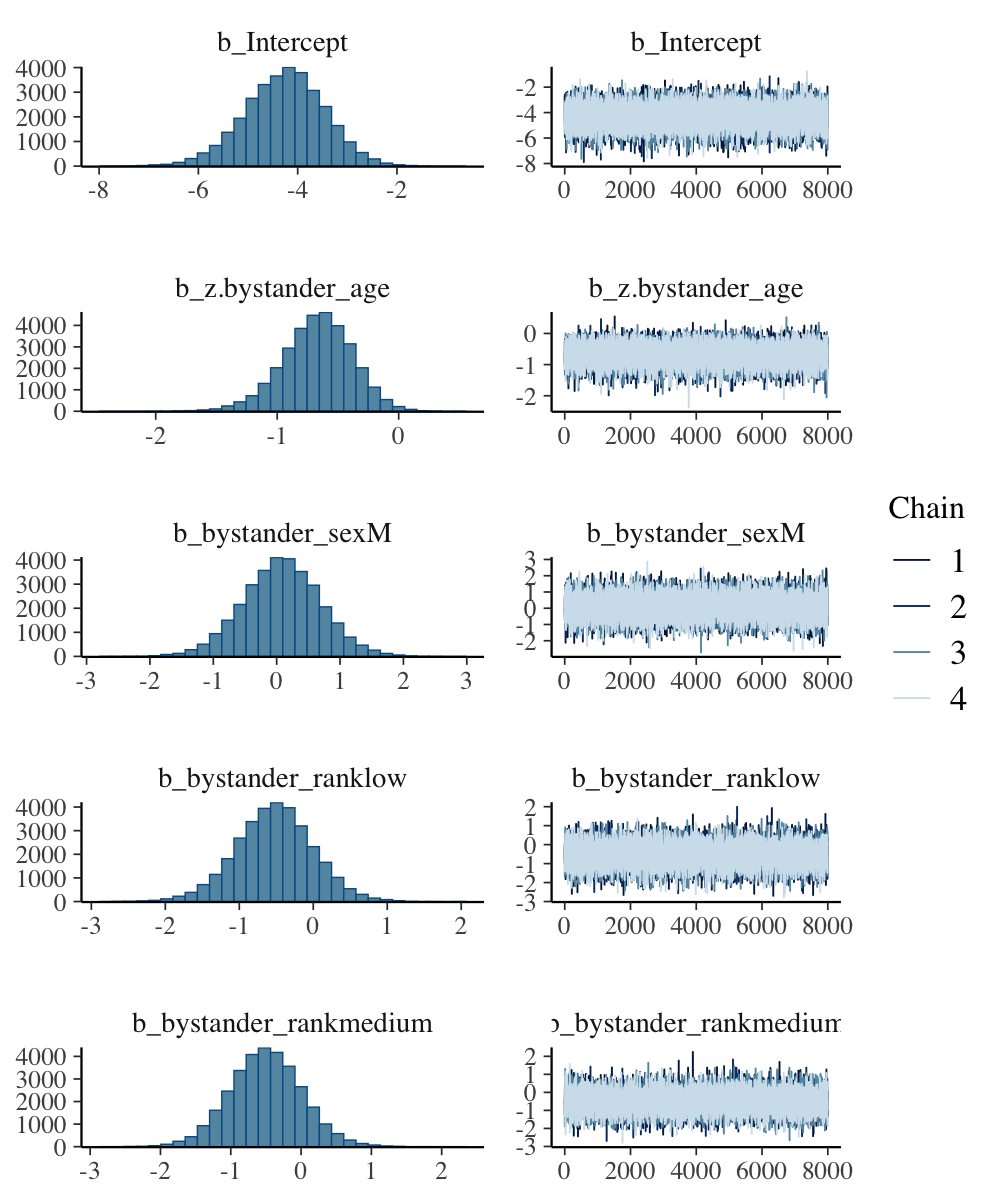

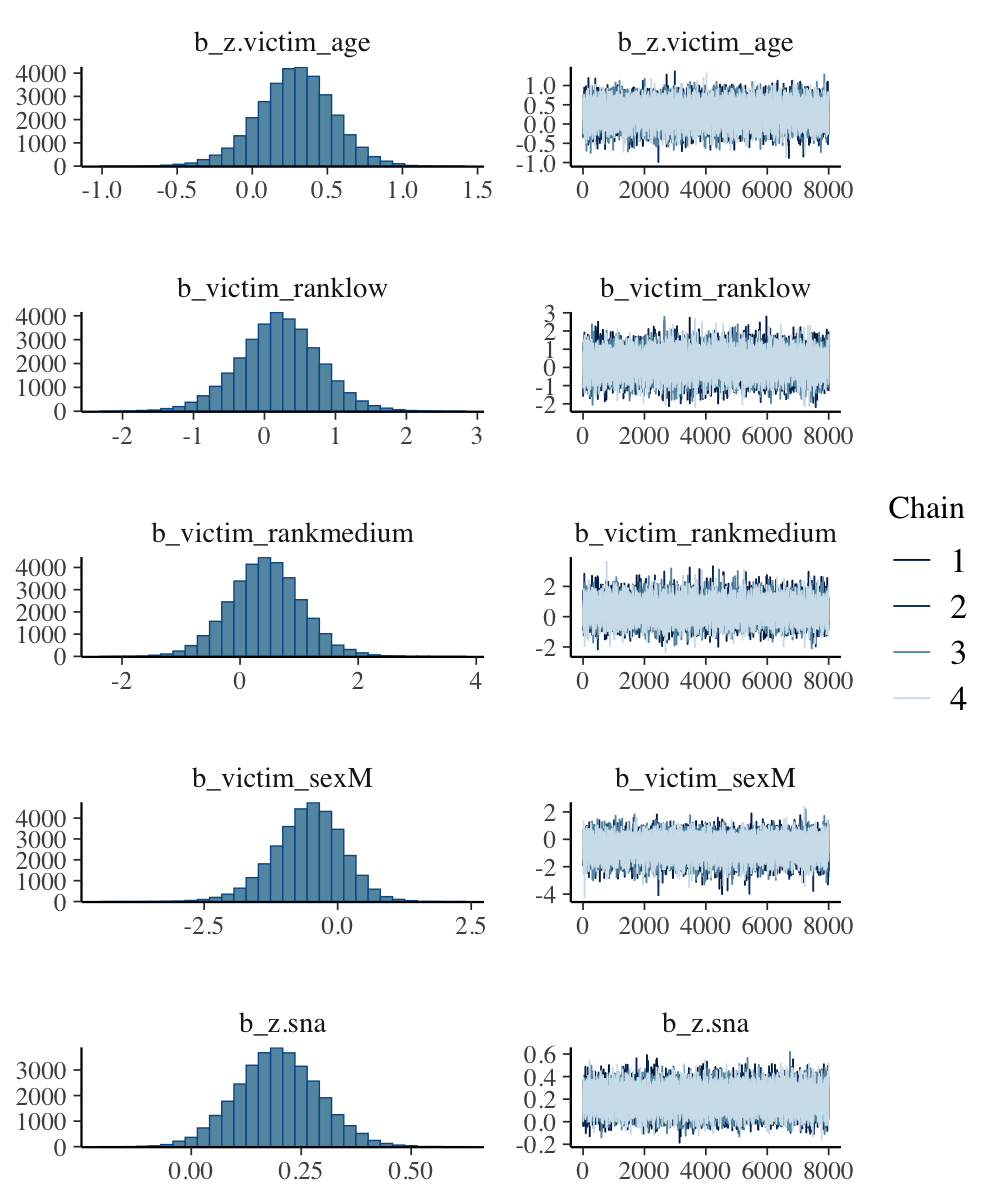

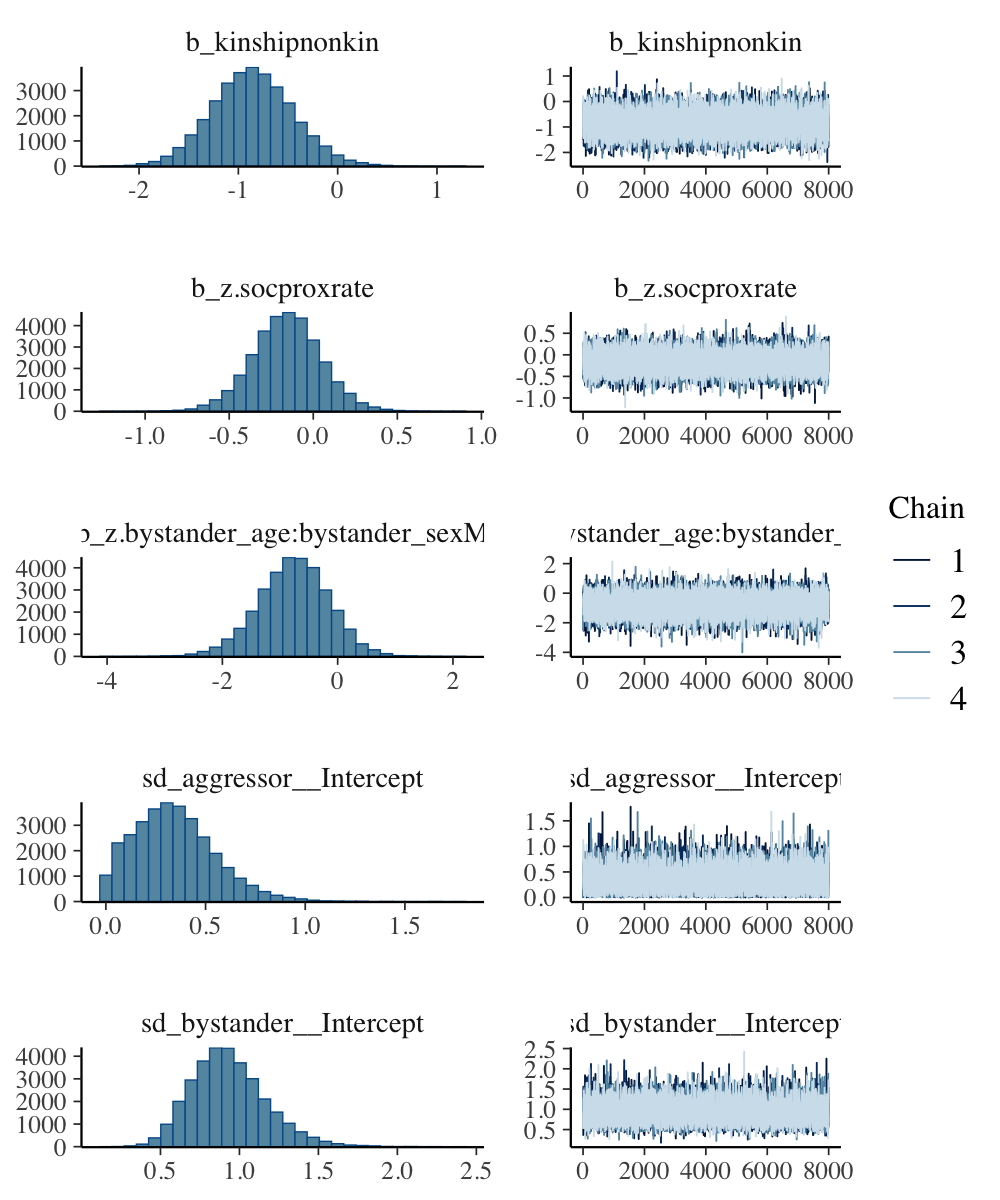

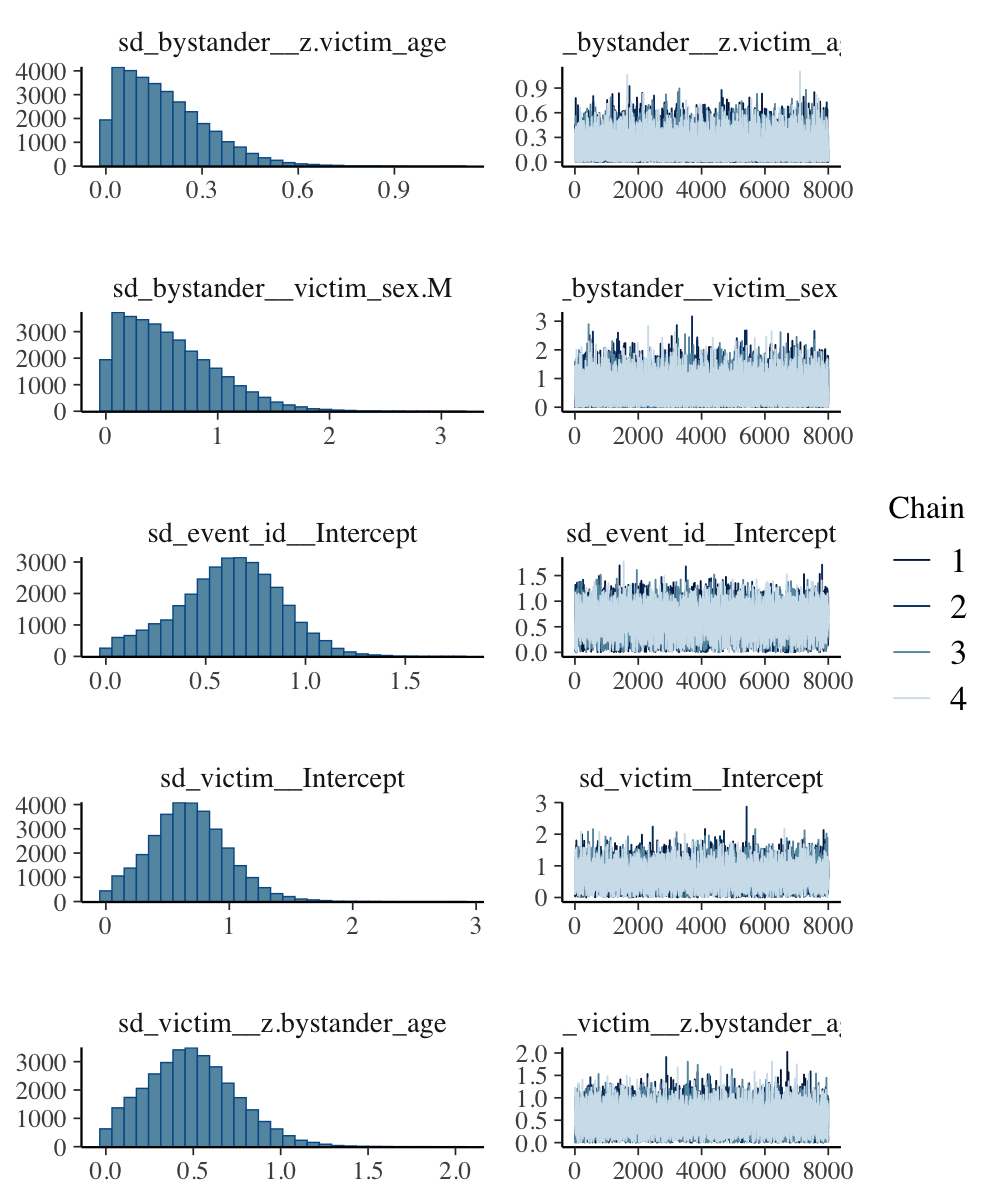

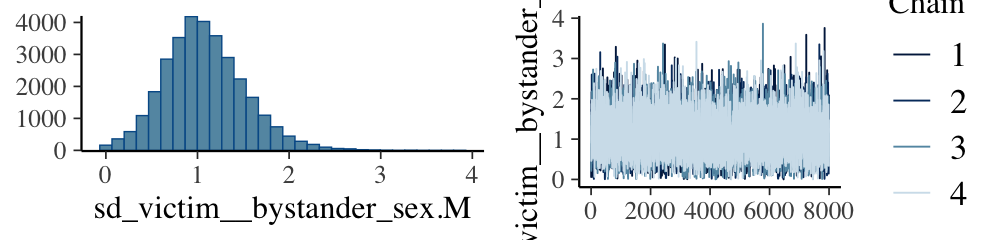

**Fig S2**. Density estimates of the observed, empirical data set y (black curves), with density estimates for 1000 simulated data sets y^rep^ drawn from the posterior predictive distribution (bright blue curves), for all models [5].

Model 1.1

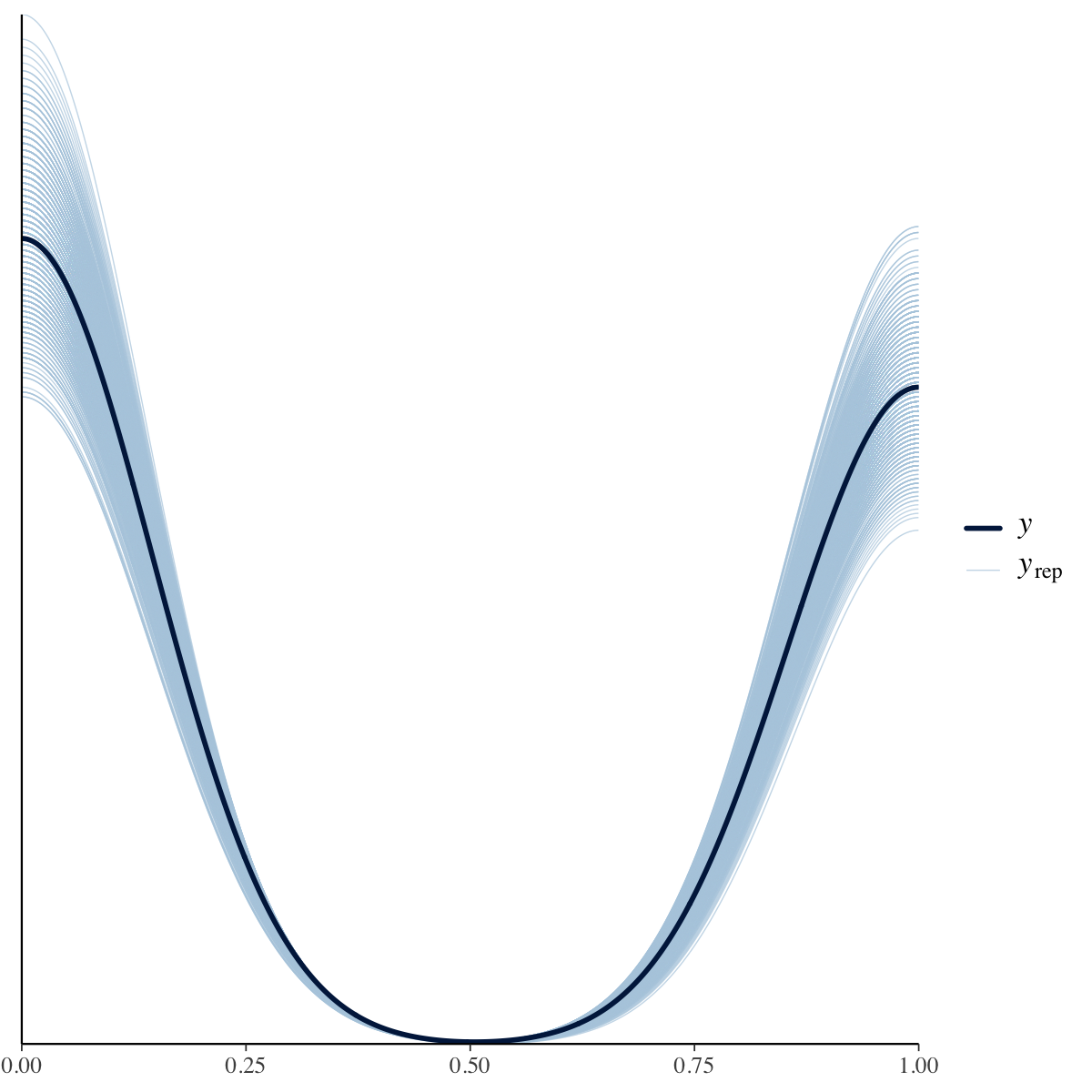

Model 1.2

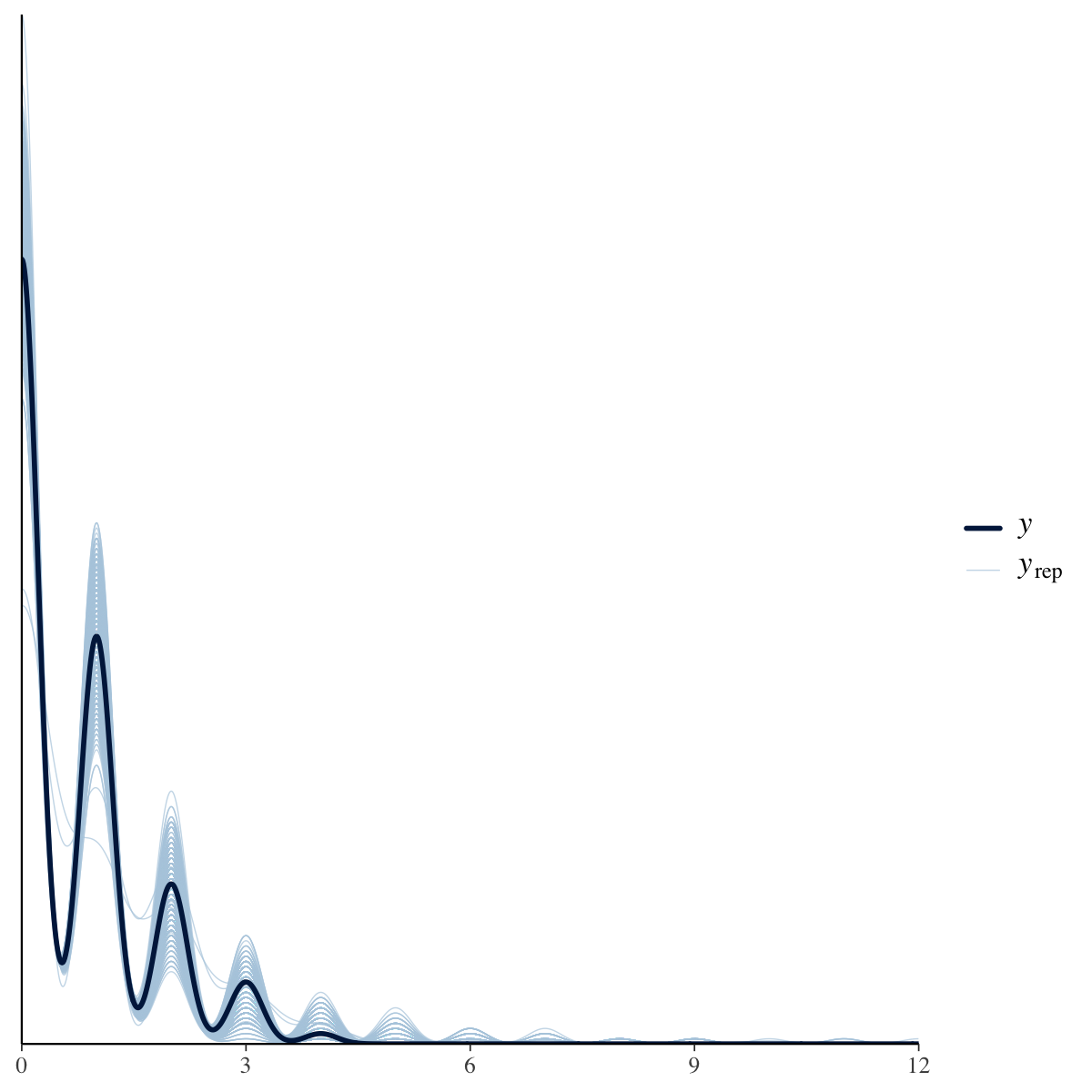

Model 2.1

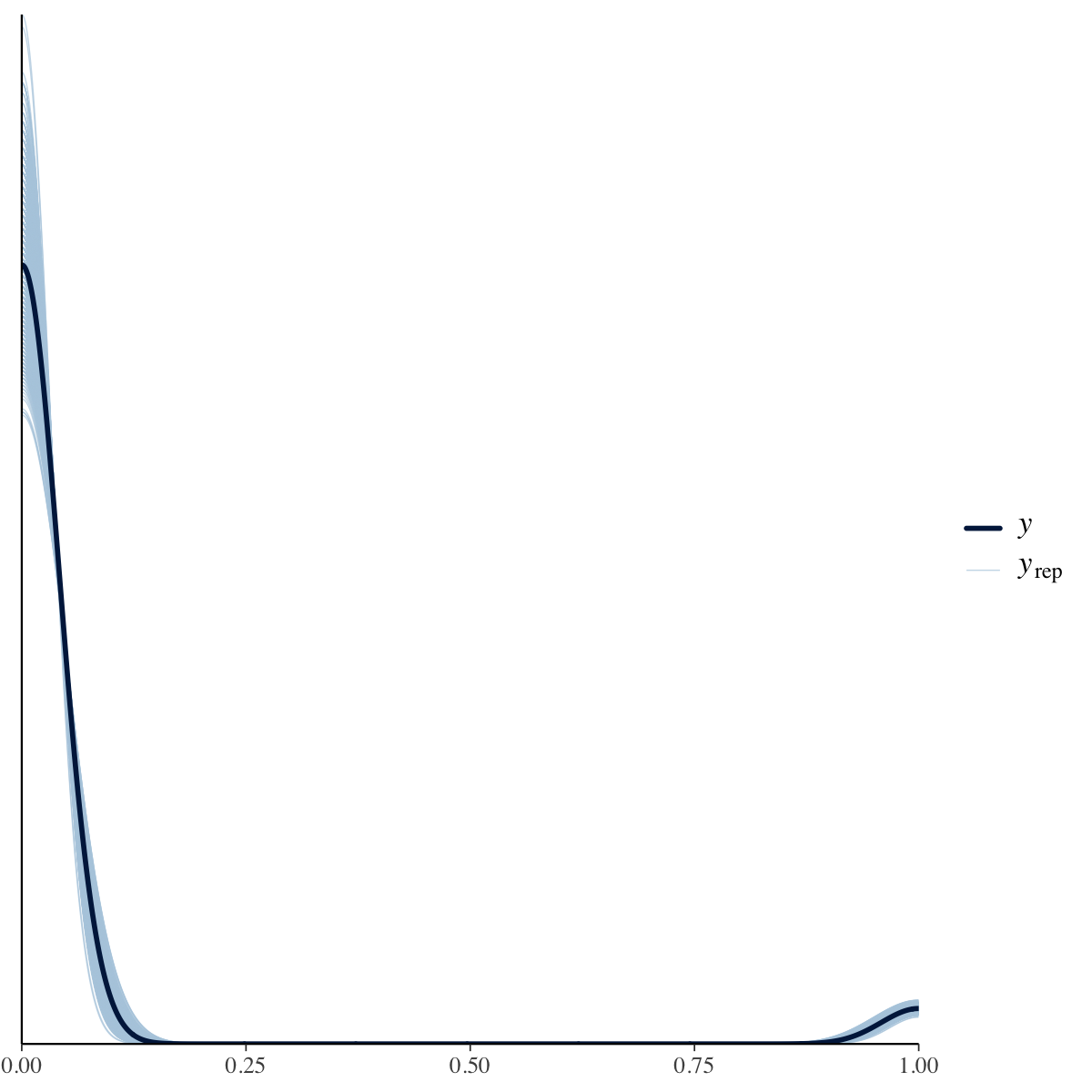

Model 2.2

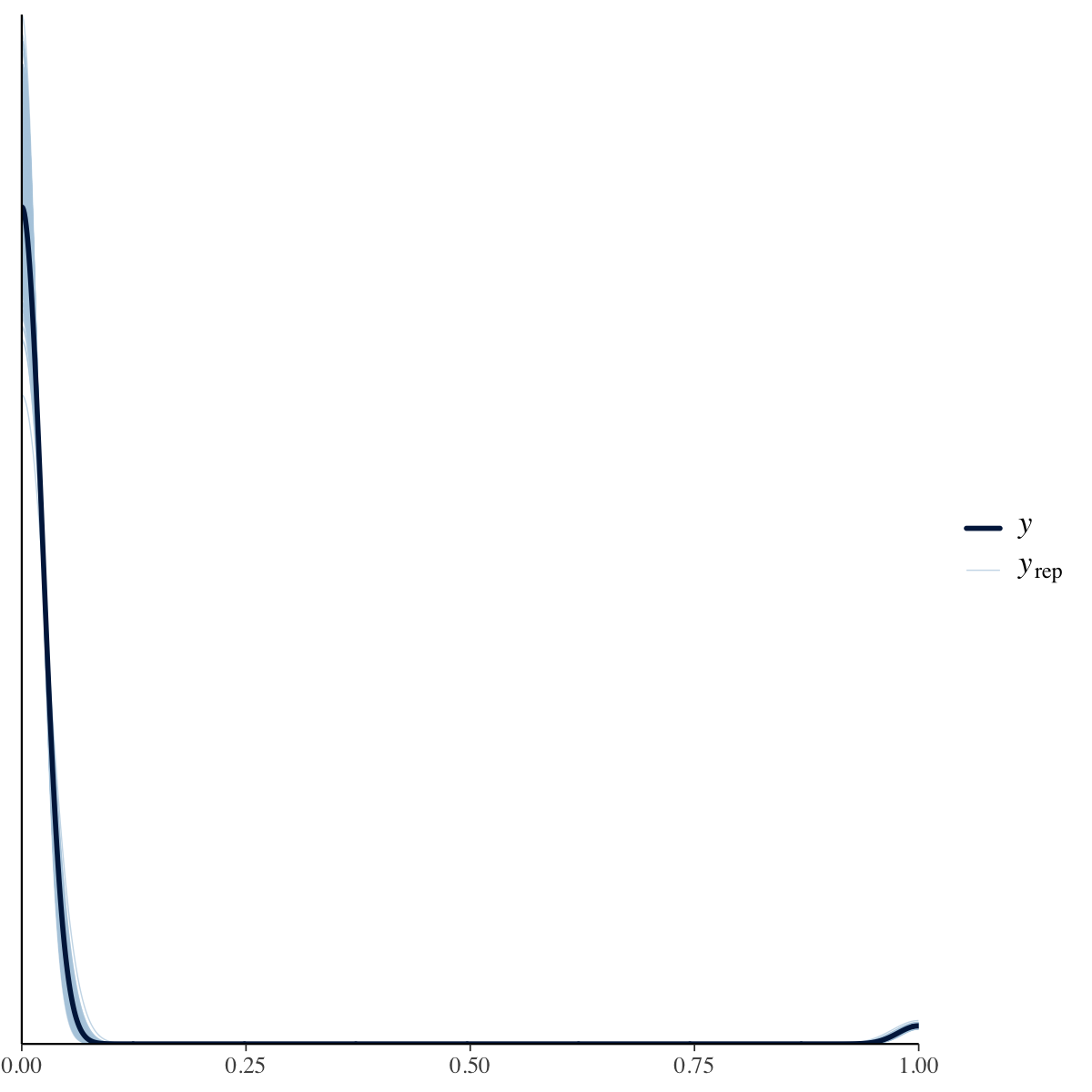

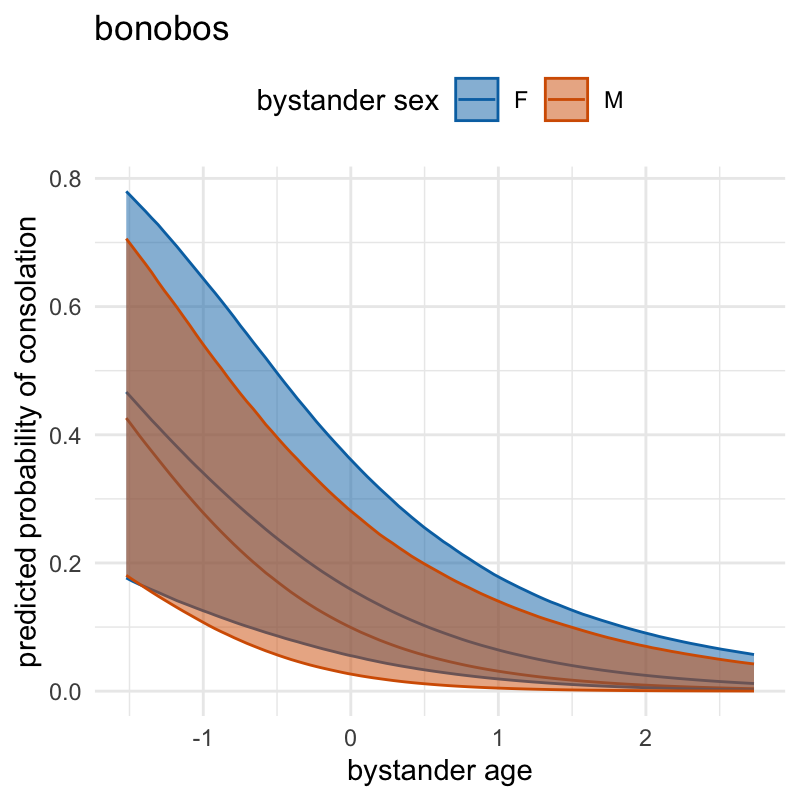

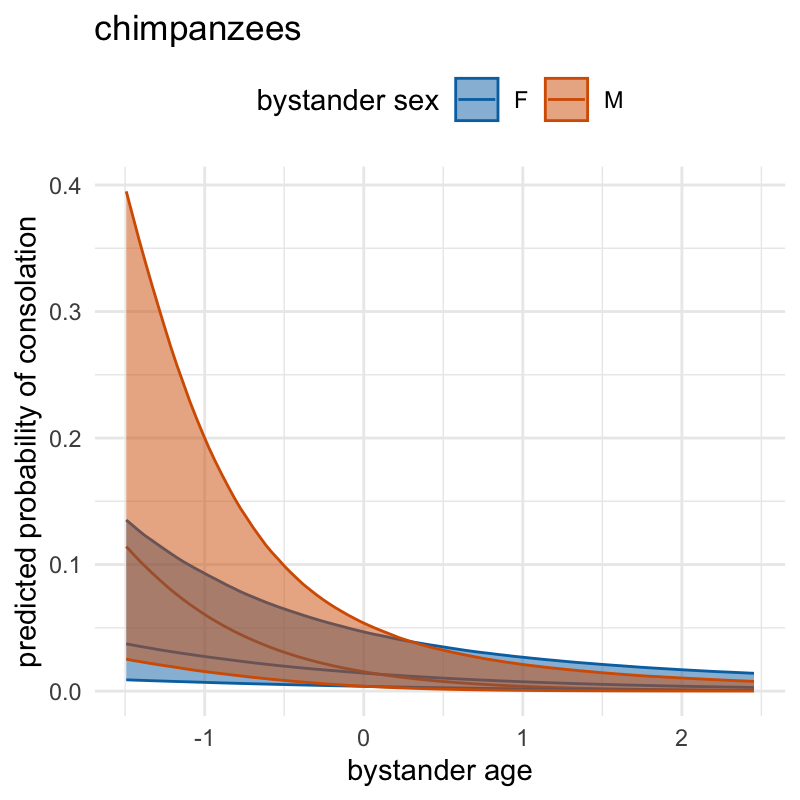
**Figure S3. Conditional probability plot for the effects of an interaction between bystander age and bystander sex and their credible intervals (89% CrI) in both species.** Note: y-axis scales are not comparable between species due to considerably more chimpanzee dyads present in the dataset.

**Supplementary data**

All data and code are provided as electronic supplementary material.
